# Genotypic and multi-environment phenotypic evaluation of the lima bean USDA National Plant Germplasm System collection

**DOI:** 10.64898/2026.06.03.729973

**Authors:** Jaclyn A. Adaskaveg, Jenna Hershberger, Andrew Farmer, R. Varma Penmetsa, Ivan Garcia-Lopez, Julian Garcia-Abadillo, Xiangming Zhou, Bao-Lam Huynh, Philip Roberts, Emmalea Ernest, Marilyn L. Warburton, Diego Jarquin, Sarah Dohle, Antonia Palkovic, Travis A. Parker, Paul Gepts, Christine H. Diepenbrock

## Abstract

Lima bean (*Phaseolus lunatus* L.) is an economically and agronomically important grain legume. Lima beans (or limas) show a range of climatic adaptations with independent domestications in the Andes (large-seeded) and Mesoamerica (small- or medium-seeded). We generated and integrated genotypic and comprehensive field- and laboratory-based phenotypic information for the available accessions in the USDA National Plant Germplasm System collection across multiple environments to inform germplasm utilization in breeding. A total of 810 accessions were genotyped using short-read, low-coverage sequencing. Accession geographic origin and domestication explained population structure. A partially overlapping subset of the panel (*n*=141-308) was field-evaluated across two years in each of Davis, CA, Central Ferry, WA, and Coachella Valley, CA (the latter was fall-planted for evaluation of photoperiod-sensitive accessions) to assess trait performance in contrasting environments. Agronomic traits such as determinacy and flowering time, and seed traits such as seed coat color and hundred-seed weight, were scored. Macronutrient traits (protein, starch, fat, and ash content) were measured on dry (mature) harvested grain via near-infrared spectroscopy. Genome-wide association analyses identified loci significantly associated with descriptive, agronomic, and seed traits, including orthologs of known genes in common bean and novel candidate regions. Genomic predictive abilities were moderate to high for key traits. Finally, we established a conditional core collection that was constrained to include 211 extensively phenotyped accessions and for which 91 supplemental accessions were selected to maximize genetic diversity from among the genotyped accessions. Overall, these resources provide a foundation to support genomics-assisted breeding of limas.

## Introduction

Dry beans are a staple crop worldwide. Among the five domesticated *Phaseolus* species, *P. lunatus* L. (lima bean) is the second most economically and agronomically important species following *P. vulgaris* L. (common bean) (Gepts 2014).

Wild lima beans originated in at least four major gene pools based on previous genomic studies (Chacón-Sánchez and Martínez-Castillo 2017; Garcia et al. 2021), including two Mesoamerican (MI and MII) gene pools associated with central-western Mexico and southern Mexico-Central America, respectively, and two Andean gene pools (AI and AII) located across Northern Peru-Ecuador and the Andes in central Colombia, respectively. At least two distinct domestications occurred, one from each of the MI and AI gene pools. Mesoamerican varieties are associated with smaller (“baby”) seeds with either round (Potato) or flat (Sieva) shapes, whereas Andean varieties are associated with large flat beans. The wide distribution of wild lima accessions in Central and South America, from Mexico to Argentina (Mackie 1943; Gutiérrez Salgado et al. 1995), suggests a relatively broad (among *Phaseolus* spp.) adaptation to climates that are generally warm and range from dry to wet (Delgado and Gama López 2015).

Approximately 90% of major lima bean germplasm collections are estimated to be photoperiod-sensitive (Dohle 2017), i.e., will only flower under short-day, long-night conditions. Photoperiod sensitivity has limited the evaluation of diverse lima accessions in US cultivation conditions (long-day summers) for priority traits expressed at or after flowering and represents a bottleneck to the utilization of these diverse accessions in breeding.

Both baby- and large-seeded limas, each with bush and vine growth habits, are bred and cultivated in multiple regions of the US. California growers produce dry limas (harvested from mature pods) for dry-packing, canning, or export (Long et al. 2014), and growers in the mid-Atlantic and Midwest (among other regions) produce succulent limas primarily for frozen or canned consumption (USDA-NASS 2019; Ernest 2020). While substantial variation in seed coat color/patterning exists among domesticated types, white-seeded dry types and green/yellow-seeded or phenolic-patterned succulent types are most commonly grown in the US. Alongside growth habit and seed coat color and size, priority agronomic traits in the US include seed yield, visual seed quality, tolerance to high temperatures (large-seeded limas are particularly susceptible to pre- and post-pollination heat stress) (Ernest 2020; Ernest and Wisser 2024), and resistance to root knot nematodes (Traverso et al. 2024) and major insect pests (such as *Lygus hesperus*; Gibson 2022). Studies of lima bean seed macronutrient composition have reported a range of protein (8.61 to 26.02%), carbohydrate (50.44 to 77.39%), fat (0.21 to 3.12%), and ash (2.35 to 6.4%) content depending on the accession, developmental stage at harvest, and growing and processing conditions (Bonita et al. 2019; Adebo 2023). Stably high protein content is of interest alongside favorable agronomic properties and maintenance of seed size- and color-based market classes to bolster the contribution of limas in human diets.

Lima beans belong to the quaternary gene pool for common bean (Parker et al. 2022), with no reports of successful crosses between the two species (Porch et al. 2013), which prevents the transfer of beneficial alleles between common and lima bean via cross-pollination. Still, the genomic resources developed in common bean have been useful in studying and breeding lima bean due to the extensive synteny of the two species, with only a few major structural rearrangements in sequenced genomes (Garcia et al. 2021; Wisser et al. 2021). Recent efforts have developed more genomic resources for lima bean including genome assemblies (Garcia et al. 2021 for domesticated MI accession G27455, Wisser et al. 2021 for Mesoamerican-derived variety Bridgeton-DES4, and Duitama et al. 2025 for domesticated Andean accession G25900) and a catalog of transposable elements and whole-genome sequences for 61 domesticated and wild accessions (Lozano-Arce et al. 2023). Approximately 500 domesticated and wild accessions have also been genotyped via genotyping-by-sequencing in Chacón-Sánchez and Martínez-Castillo (2017), Garcia et al. (2021), and Gibson (2022). These resources in lima bean have allowed for further exploration of population structure and the genetic basis of important traits within the species.

Here, we investigated a panel of 810 lima bean accessions representing the available inventory (both domesticated and wild accessions) from the USDA National Plant Germplasm System (NPGS) lima bean collection complemented by other domesticated and wild accessions of relevance to lima bean breeding and genetics. The accessions were subjected to single-seed descent and genotyped using low-coverage, short-read sequencing; population structure and linkage disequilibrium decay were assessed. We field-evaluated a total of 353 accessions across six field environments: two years in each of Davis, CA, Coachella Valley, CA, and Central Ferry, WA. Agronomic traits including determinacy and flowering time were scored, and hundred-seed weight (HSW) and near-infrared spectroscopy (NIRS)-predicted macronutrient traits were assessed on dry (mature) grain harvested from these trials. To facilitate NIRS-based predictions, we developed a custom calibration for use on ground dry lima beans, which was compared to a pre-existing calibration. Then, we conducted genome-wide association studies and genomic prediction for these agronomic and seed traits, alongside descriptive traits that were scored in the genotyping process or available in the USDA-ARS Germplasm Resources Information Network (GRIN). Finally, we used a super-saturated design to identify conditional and unconstrained (Virdi et al., 2023; Persa et al., 2023) core collections from the genotyped accessions. The conditional core was constrained to include accessions that have been tractable to phenotyping in multi-environment trials based on successful seed multiplication. Overall, the data and results presented in this study provide both genotypic and phenotypic characterizations of the USDA NPGS lima bean collection to provide an integrated set of resources towards the comprehensive improvement of limas.

## Materials and Methods

### Plant material

The accessions in this study (Table S1) were primarily obtained from the USDA NPGS lima bean collection held at the Plant Germplasm Introduction and Testing Research Station in Pullman, WA (National Plant Germplasm System 2025). Specifically, all accessions from the collection that were available (i.e., had >300 seeds in inventory) and a subset of those with 200 to 300 seeds in inventory as of October 2022 were provided. Breeding materials from UC Davis, the University of Delaware, and commercially available heirloom lines were also included. A subset of accessions was sourced from a previous diversity panel (Gibson 2022) originally obtained from the International Center for Tropical Agriculture (CIAT, Cali, Colombia) or from materials in the UC Davis breeding program, such as the parents of a diallel. Accession geographic origin was integrated from USDA-ARS GRIN (https://www.ars-grin.gov/) and the Genesys-hosted CIAT database (https://www.genesys-pgr.org/) where available.

### Genotyping

Leaf tissue of one plant per accession was sampled for genotyping in a greenhouse at UC Davis, USDA Central Ferry Farm, or at the Clemson University Pee Dee Research & Education Center. Dry (mature) seed harvested from the sampled plant was retained to enable future propagation of the genotyped stock. Approximately 2-3 young, unexpanded trifoliate leaves (and part of a slightly older, larger leaf as needed) were collected from individual plants. Leaf samples were placed in 96-well plates and shipped to the HudsonAlpha Institute for Biotechnology (Huntsville, AL). A total of 1,034 samples were genotyped, 810 of which were unique accessions, and an additional 224 duplicate accessions (for which a single seed was planted at each of UC Davis and Clemson) were included for internal control. DNA extraction was performed by HudsonAlpha using bioecho EchoLUTION (BioEcho Life Sciences GmbH, Cologne, Germany). Library preparation was performed by HudsonAlpha via Twist 96-plex library preparation kit (Twist Bioscience, South San Francisco, CA). Genotyping via short-read, low-coverage sequencing was performed by HudsonAlpha using their Khufu platform (Korani et al. 2021), and the subset of samples with coverage of <0.5 were re-run on the sequencer.

### Read alignment and variant discovery

Calling of single-nucleotide polymorphisms (SNPs) was conducted by HudsonAlpha using their KhufuVAR group of software tools. Briefly, they divided the population into sets of 100 or fewer samples and ran a Get Loci Structure module to call SNPs, filtering by depth. This module recovers reads from BAM files and accounts for them at a given physical position for purposes of counting reads and calling SNPs. A minimum depth of four reads (and maximum depth of 3,000 reads) was required to call a SNP. Unimputed data were used in all downstream analyses.

Sites were filtered in TASSEL 5 (Bradbury et al. 2007) with a minimum site count as the number of taxa divided by two, minimum minor allele frequency (MAF) of 0.05, and maximum heterozygous proportion of 0.8, resulting in a total of 29,409 SNPs. Subsequent taxa filters were applied for subsets of accessions analyzed as part of the genome-wide association studies described below. For each subset of accessions analyzed, the same thresholds for minimum site count, minimum MAF, and maximum heterozygous proportion were applied based on the accessions analyzed (Figure S1).

### Analysis of population structure

To investigate population structure, multiple approaches were compared using the 29,409 filtered SNPs. To visualize genetic structure and variation across accessions, a principal component analysis was conducted in TASSEL and visualized in R. The variational Bayes inference algorithm fastSTRUCTURE (Raj et al. 2014) was used to estimate structure with a large dataset. Successive numbers of subpopulation (K) values from 2 to 20 were tested using default options. The optimal K was determined and reported using the *chooseK.py* function. K=3 informed the major subpopulation groups identified in this study. Neighbor-joining tree analysis was performed using the Cladogram function in TASSEL with ‘Neighbor_Joining’ selected as the clustering method and visualized with the *ape* package in R (Paradis and Schliep 2019). Accessions were categorized into major subpopulation groups based on the fastSTRUCTURE, PCA, and neighbor-joining tree analysis and overlaid with accession origin, improvement status, and hundred-seed weight from USDA-ARS GRIN.

### Linkage disequilibrium

Linkage disequilibrium (LD) was estimated in *Plink* v1.90b7.1 (www.cog-genomics.org/plink/1.9/; Chang et al. 2015) with specifications of –r2, –allow-extra-chr, and –make-founders, and with – ld-window-r2 set to 0. Specifically, using the filtered datasets described above, LD was estimated (1) for the full set of genotyped accessions included in this study (*n*=1,034); (2) for the subpopulations determined from fastSTRUCTURE analysis (Andean (*n*=213), Mesoamerican (*n*=629), *k6* and *k7* (*n*=86), and admixed (*n*=192)); and (3) on each of the input data sets used for GWAS, including the 619 Mesoamerican accessions that were scored for descriptive traits, the 248 Mesoamerican accessions evaluated in multi-environment trials, and the 127 Mesoamerican accessions evaluated for flowering time in Davis 2024 and 2025 (Figure S2). The resulting output, in .ld format, was plotted using the plot() and lines(lowess()) functions in base R (R Core Team 2024) via RStudio Server (Posit Team 2025).

### Field trials

A subset of the genotyped panel was field-evaluated in three locations in each of two years. The seed stock derived from the genotyped plant, or from subsequent greenhouse- or field-based multiplications of that stock, was used for field planting.

In summers 2023 and 2024, a subset of day-neutral (photoperiod-insensitive) accessions was grown in Central Ferry, WA (46.6711, -117.7540). 118 accessions were planted on May 16, 2023, by hand. Half of each plot was direct-seeded, and half were one-month old transplants, as it was uncertain whether the growing season would be long enough to produce mature seed. A completely randomized design was used with one replicate. Plots were 1.5 m in length with 10 plants per plot, and 1.5 m of spacing was included between plots. In summer 2024, 99 accessions including 89 of the same day-neutral accessions grown in 2023 were planted by hand on May 16, 2024, with the same spacing. Two replicate plots per accession were included in an augmented incomplete block design using the determinate cultivars UC Beija-Flor and UC 92 as checks. No supplemental fertilizer was used as soil tests showed normal levels of fertility. Buried drip was used for irrigation both years. Plots were hand-harvested in both years throughout October.

In fall 2023 and 2024, photoperiod-sensitive and day-neutral accessions were grown in UC Riverside’s Coachella Valley Agricultural Research Station (CVARS) located in Thermal, CA (33.5201, -116.1515). In 2023, 247 accessions were hand-planted on November 1 in an augmented incomplete block design with up to two replicate plots per accession as seed availability allowed. Plots with length of 2.1 m were sown with 30 seeds, and plots were spaced 0.9 m apart and planted in every other bed; the beds were 0.9 m wide, and drip irrigation was used. Checks UC 92 and UC Beija-Flor were included in each block. In both years, 544 kg of 15-15-15 starter fertilizer was applied, resulting in 201 kg per ha of nitrogen. In addition, 454 kg of gypsum was applied. During the season of both years, 94 L per ha of urea ammonium nitrate (UAN) 32 and 56 kg per ha of 20-20-20 fertilizer was applied, resulting in a total of 15 kg per ha of nitrogen. Plots were hand-harvested between May 6 and 28, 2024. On September 15, 2024, 258 accessions were hand-planted in an augmented incomplete block design with the same checks. Accessions were separated into two super-blocks based on growth habit. Bush-type accessions were planted on every bed in one block, while vine-type accessions were planted on every other bed. The bush super-block was hand-harvested between February 10 and 19, 2025. The vine super-block was hand-harvested between March 11 and 31, 2025.

In summers 2024 and 2025, day-neutral accessions were grown in Davis, CA (38.5366, - 121.7772). In 2024, 178 accessions were planted on May 23 and 24 via self-propelled Wintersteiger planter (Ried im Innkreis, Austria) in 1.8 m plots sown with 30 seeds; certain large-seeded accessions were hand-planted. Up to two replicate plots were planted in an augmented incomplete block design using UC 92, UC Beija-Flor, and Jackson Wonder as checks. In 2025, 186 accessions were planted on May 22 via self-propelled Wintersteiger planter; certain large-seeded accessions were hand-planted. In each of these two years, starter 8-24-6 fertilizer was applied at a rate of 281 L per ha, resulting in 31 kg per ha of nitrogen. Beds were 1.5 m center to center and irrigated via buried drip. Plants were cut and windrowed, and plots were harvested with a ZÜRN 150 (Westernhausen, Germany) small-plot research combine on October 16, 17, and 21, 2024, and October 7–8, 2025.

### Descriptive traits and hundred-seed weight (HSW) phenotyping

The weight of 100 seeds was measured for dry (mature) seed harvested from all field environments. When fewer than 100 seeds were available, seeds were counted and total weight was measured, then HSW was calculated by averaging the seed weight and multiplying by 100. In Davis 2024 and 2025, flowering time was recorded when 50% of plants in a plot had at least one open flower. Seed coat color was categorized into pigmented or white/green based on the initial seed used for single-seed descent. Plots were categorized in the Davis and CVARS fields as vine or bush types based on indeterminate or determinate growth, respectively. Flower color was scored as pigmented or white in the greenhouse and in the Davis and CVARS fields. Photoperiod sensitivity (sensitive or day-neutral) was based on USDA-ARS GRIN data, with re-classifications to ‘sensitive’ made (as reflected in Table S1) for a small number of accessions that were annotated as day-neutral in GRIN but did not flower in the Davis or Central Ferry environments.

### Nutritional trait phenotyping

Grain macronutrient content for protein, starch, fat, and ash (total minerals) was quantified as a percentage of grain composition using a benchtop near-infrared spectroscopy (NIRS) instrument (DS2500; FOSS, Eden Prairie, MN, USA). In preparation, at least 6 g of mature dry seed per sample were ground via IKA Tube Mill 100 (IKA-Werke GmbH & Co. KG, Staufen, Germany) at 12,000 rpm for 1 min and 20 s in intervals of 30 s. Ground powder was stored at -20°C until analysis and brought to room temperature before total mass was emptied into a FOSS small sample cup. Spectral data were acquired over the wavelength range of 400 to 2500 nm at 0.5 nm intervals and recorded in log (1 R^-1^) format (ISIscan NOVA) with the pre-existing Vegetal Protein Meals calibration selected (FOSS S800444, Hillirød, Denmark), which generated macronutrient predictions for each sample. Raw spectral data were exported for development of custom calibrations as described below.

A subset of samples were selected for reference wet-chemistry analysis. A total of 320 samples, including 40 samples from each of the six field environments tested and 80 additional lima samples across multiple environments, were selected via Kennard-Stone algorithm to identify a uniform distribution of samples based on NIR spectral data. Sample selection was performed using the “kenStone()” function in the R package *prospectr* (Stevens and Ramirez-Lopez 2025) with metric set to ‘mahal’ and pc=0.99. Samples were submitted to the UC Davis Analytical Lab for protein (AOAC Official Method 990.03), fat (AOAC Official Method 2003.05), and ash (AOAC Official Method 942.05) content, and moisture (calculated from dry matter). Reference wet-chemistry data for total starch was generated on the same 40 samples from each of the six environments via Megazyme (Neogen Corporation, Lansing, MI, USA) Total Starch (α-amylase/amyloglucosidase) assay protocol kit (AOAC Method 996.11).

A custom NIRS calibration was developed using the wet-chemistry data as reference values using partial least squares regression (PLSR) via the *pls* package in R (Liland et al. 2026). For each trait, five iterations of five-fold cross validation with random assignment of samples to folds were conducted with a maximum of 20 components. The number of principal components used for each trait was selected based on root mean squared error of prediction (RMSEP) using the ‘selectNcomp’ function with the ‘onesigma’ method (Table S2). Multiple pre-treatments (Savitzky-Golay first and second derivatives, and standard normal variate (SNV)) were evaluated and compared to no pre-treatment. After evaluating RMSEP and the correlation between predicted and observed values for each trait on samples analyzed via wet chemistry, no pre-treatment was selected, and predictions were then generated for the full sample set using the ‘predict’ function.

Macronutrient traits are reported as a percentage of grain composition. Given variability of moisture between samples and across environments, protein, starch, fat, and ash content were adjusted for moisture content by dividing the trait of interest by the dry matter and multiplying by 100, calculated with the following formula: (Trait / (100 - Moisture))*100

Predicted values from the custom calibration and from the pre-existing Vegetal Protein Meals calibration were compared against wet-chemistry reference values, and the Pearson correlation coefficient (*r*) was calculated and reported (Table S2). The custom calibration-based predictions were then utilized in downstream analyses.

### Analysis of phenotypic data

Phenotypic transformations were applied using Box-Cox transformations to better meet the assumption of normality of residuals for subsequent genome-wide association studies (GWAS). Pearson’s correlations were calculated to examine trait relationships. The main and interaction effects of genotype and environment for traits assessed across multiple environments (flowering time, HSW, and grain macronutrient content) were analyzed using the *statgenGxE* package (Van Rossum 2025) in R using the createVarComp() function and a custom modification of the vc() function as described in LaPorte et al. (2026). The multi-environment analysis across all trials was implemented with the following model: *Yijkl=μ+Li+T(L)ij+Gk+(G×L)ik+(G×T(L))ijk+R(T(L))ijl+εijkl* where *Yijkl* is the observed phenotype, μ is the overall mean, *L*i is the fixed effect of location i, *T(L)ij* is the random effect of year *j* nested within location *i*, *Gk* is the random effect of genotype *k*, *(G×L)ik* is the random interaction between genotype and location, *(G×T(L))ij* is the random interaction between genotype and year nested within location, *R(T(L))ijl* is the random effect of replicate *l* within each location-year, and ε*ijkl* is the residual error term. Analysis within a location was performed with the following model: *Yijkl=μ+Ti+Gj+(G×T)ij+R(T)ik+εijkl* where *Yijkl* is the observed phenotype, μ is the overall mean, *Ti* is the fixed effect of year *i*, *Gj* is the random effect of genotype *j*, *(G×T)ij* is the random interaction between genotype and year, and *R(T)ik* is the random effect of replicate *k* within year.

### GWAS

For each trait, GWAS was conducted using the filtered genotypic data described above for accessions with phenotypic data. Three sample sets were tested in GWAS: (1) accessions within the Mesoamerican group determined from fastSTRUCTURE analysis with descriptive traits characterized by greenhouse growout or through data obtained from USDA-ARS GRIN or CIAT (*n*=619), (2) a union of accessions grown across the six environments within the Mesoamerican group defined from fastSTRUCTURE analysis to be *k1*, *k3*, *k4*, *k5*, and *k8* (*n*=248), and (3) a union of accessions characterized for flowering traits in the Davis 2024 and Davis 2025 environments within the Mesoamerican group (*n*=127). For each subset of accessions analyzed, genotypic data was filtered using a minimum site count as the number of taxa divided by two, MAF=0.05, and maximum heterozygosity=0.8. A total of (1) 29,409, (2) 19,398, and (3) 13,178 SNPs were utilized for each respective sample set. In each analysis, multiple models were compared including a mixed linear model (MLM), FarmCPU, and BLINK using kinship and two principal components as covariates to account for population structure. MLM and FarmCPU were performed using the *rMVP* package (Yin et al. 2026) and BLINK using the *BLINK* package (Zhang and Zhou 2025) in R. Kinship and principal components were generated with *rMVP* for each subset of accessions. Manhattan and Q-Q plots were generated from the output in R using *CMplot* (Lin-Yin 2024). Significant associations were reported using Bonferroni adjustment with α=0.05.

Candidate genes were identified in the *P. lunatus* G27455 reference genome (Garcia et al. 2021) in the Legume Information System (LIS; https://www.legumeinfo.org/) (Berendzen et al. 2021). The candidate gene search space for each GWAS dataset was determined based on the physical distance by which LD decayed to *r* = 0.2, corresponding to 650 kb for the descriptive trait dataset and 500 kb for the multi-environment seed trait dataset (Figure S2). Significant hits that were identified in multiple methods and/or environments were mined for genes and gene descriptions within the search space using LegumeMine v. 5.1.0.3. Then, synteny with common bean (*P. vulgaris*) was identified using LIS Pangenome lookup. Gene expression in the relevant tissue was evaluated using *P. vulgaris* Gene Expression Atlas (O’Rourke et al. 2014). Results with synteny to common bean and gene expression in common bean were then examined for key words related to the trait of interest. Gene functional annotations such as GO terms and InterPro were examined for candidates. Homology was further examined for identified candidates via BLASTp (Camacho et al. 2009) using the respective lima amino acid sequence in FASTA format from LIS.

### Genomic prediction

The GBLUP method was implemented for genomic prediction (GP) of descriptive, agronomic, and seed traits using the TASSEL 5 Genomic Selection plugin (Bradbury et al., 2007; LaPorte et al., 2024). A kinship matrix was created from each SNP dataset used in GWAS (described above) using the TASSEL kinship matrix function. GBLUP was conducted using 20 replicates of a five-fold cross-validation, a typical framework for GP, for each trait in each environment.

### Selection of conditional and unconstrained core collections

The starting set for this analysis was the 180 accessions that had already been evaluated in four or more environments within the broader project of which this work was part as of March 2025, plus 36 accessions designated as photoperiod-sensitive in USDA-ARS GRIN that had been phenotyped in three or more environments as of March 2025. The set of environments considered in these counts were five of the six field environments studied herein (all except for Davis 2025, which had not yet taken place), and 2024 evaluations in two additional field locations (at Clemson and a separate nematode resistance screen at CVARS; manuscripts in preparation). Those accessions summed to a total of 305 samples, if counting duplicates; i.e., accessions that underwent separate single-seed descent and genotyping by both the UC Davis and Clemson teams. These 305 samples were constrained to be in the conditional core collection. A set of 100 supplemental samples was then selected from among the 1,034 genotyped samples (if counting duplicates) to maximize genetic diversity using a super-saturated method via parallel computing as described in Virdi et al. (2023). Briefly, this method aimed to identify the sample set of (305 + 100) samples that minimized E(S^2^), where E(S^2^) is the sum of the squared values of the diagonal elements of the genetic similarity matrix (S) between pairs of SNPs. First, a set of 100 supplemental samples was randomly selected. E(S^2^) was calculated on that set, and one sample from the unselected cohort (of 629 samples) was then swapped into each of the 100 positions. This procedure was continued until all samples had been swapped in (along both the rows and the columns), resulting in a large number of estimates of E(S^2^) (namely, 100 x 629) for each iteration of running the method. The optimized subset was then used as the starting point for the next iteration, and iterations continued until the minimum E(S^2^) from the previous (k^th^) iteration was less than or equal to that of the current (k+1)^th^ iteration.

The same super-saturated method was also run using the 305 samples constrained to be in the conditional core, but then selecting a set of 100 supplemental samples (from the 1,034 samples) that minimized genetic diversity (i.e., maximized E(S^2^)) to generate a recommended initial training set for future use in genomic prediction/selection.

Finally, the same super-saturated method was used to select an unconstrained core of 402 samples from among the 1,034 samples. This unconstrained core was designed to be similar in size to the conditional core (counting constrained plus supplemental samples), to enable determination of the extent of genetic diversity captured by the conditional vs. unconstrained cores.

## Results

### Genotyping

Genotypic data were returned for a total of 1,034 samples, representing 810 unique accessions. A total of 134,309 SNPs were present in the raw data set prior to filtering (Figure S1). Of these 810 accessions, 224 accessions were taken independently through single-seed descent and genotyped at each of UC Davis and Clemson. These duplicate samples within the genotypic data were generally proximal to each other in principal component space with limited exceptions (Figure S3).

### Population structure and genetic diversity

The panel of 810 accessions was sourced from 37 countries (five continents) (Fig. 1a) and included wild and domesticated accessions. A filtered set of 29,409 SNPs was utilized for population structure analyses. The fastSTRUCTURE analysis indicated eight groups (K=8) could explain population structure (Fig. 1b, Figure S4). At K=8, 81.43% of accessions were assigned to a group with ancestry greater than 70%. The remaining 192 accessions with less than 70% ancestry were not assigned to a group and were categorized as admixed. The group membership was largely explained by seed size, continent of origin, and domestication status (Table S1, Table S3). For example, large-seeded limas, associated with Andean origins, were primarily assigned to *k2* (average HSW based on USDA-ARS GRIN = 94 g) and included accessions primarily from Peru, Ecuador, and the US. *k7* was primarily composed of domesticated accessions from Brazil and had a wide range of seed sizes (average HSW = 66 g, range = 31-127 g). *k6* and *k7* were grouped together at other values of K; however, unlike *k7*, *k6* was primarily composed of wild accessions from Central America, including Guatemala and Costa Rica (average HSW = 10 g). Baby-seeded domesticated limas, associated with Mesoamerican origins, made up the remaining clusters (*k1*, *k3*–*k5*, and *k8*; range of average HSWs = 31–48 g, Table S3). Group membership corresponded to continent of origin and improvement status (Table S3, Figure S4). Notably, *k5* and *k8* primarily represented US accessions, *k4* primarily represented Brazilian (and secondarily, Colombian) accessions, while *k1* and *k3* primarily represented Central American (Guatemala and El Salvador) and Mexican accessions, respectively.

**Fig. 1.**
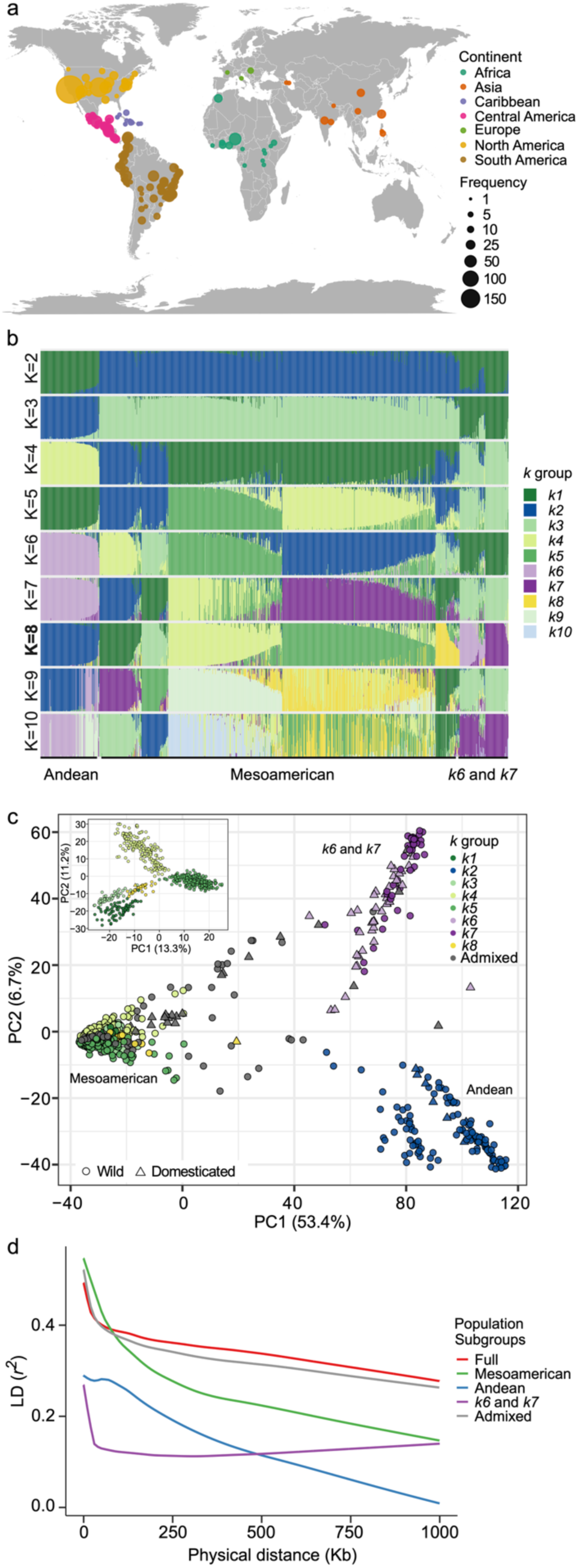
Characterization of the USDA NPGS *Phaseolus lunatus* panel. **a)** Global distribution of the panel based on the level of information available on geographic origin. The panel was grouped into population subgroups using **b)** fastSTRUCTURE (K=1–10) analysis, with K=8 being the optimum, **c)** principal component analysis (PCA) of the full panel and inset of PCA of leftmost (Mesoamerican) accessions, each colored by *k* group at K=8 and with shapes indicating wild or domesticated, **d)** linkage disequilibrium for the full panel and population subgroups. The population subgroups were defined as Mesoamerican (*k1, k3, k4, k5, k8*), Andean (*k2*), *k6 and k7*, and Admixed (accessions with <0.7 identity to a *k* group).

A principal component analysis (PCA) of the genotypic data showed that three PCs explained 62.2% of the variation in the panel. PC1 explained 53.4% of the variation and largely separated subgroups *k2*, *k6* and *k7* from the other *k* groups. The second and third PCs explained 6.7% and 2.6% of the variation, respectively. PC3 explained variation among the small-seeded accessions in *k1*, *k3*–*k5*, and *k8* (Figure S4); a PCA run on the samples in this cluster revealed *k4* and *k5* were separated from *k1*, *k3*, and *k8* across both PC1 and PC2 (Fig. 1c inset). Based on these fastSTRUCTURE and PCA results, we identified three subpopulations: a Mesoamerican group (*k1*, *k3*, *k4*, *k5*, *k8*), an Andean group (*k2*), and a group with mixed origins (*k6* and *k7*; Fig 1c). *k*2 remained distinct from all other *k* groups in a neighbor-joining tree analysis, while *k6* and *k7* clustered together and were related to *k1* (Figure S4).

Linkage disequilibrium (LD) decay was evaluated for the full panel, the subpopulations defined above, and the admixed individuals (Fig. 1d). The full panel decayed to an *r*^2^ value of 0.3 near 1,000 kb, and the admixed group displayed a similar pattern to the full panel. The Mesoamerican group reached an *r*^2^ of 0.2 by 700 kb, while the Andean group reached an *r*^2^ of 0.2 by 200 kb. In contrast, the *k6* and *k7* groups displayed low *r*^2^ values at short distances (Figure S5).

### Descriptive traits across the panel

Accession-level descriptive characteristics, independent of growing environment, were determined across the panel including seed coat color (white or pigmented), determinacy (determinate or indeterminate), flower color (white or pigmented), and photoperiod sensitivity (sensitive or day-neutral) (Table S4). No pairwise combination of these traits had a Pearson’s |*r|* > 0.3 in a correlation test of all accessions (Table S5). Correlations for the subpopulations determined in population structure analysis were also analyzed. Mesoamerican accessions showed significant correlations between determinacy and photoperiod sensitivity (*r* = -0.540) and between seed coat color and photoperiod sensitivity (*r* = 0.480), corresponding to the higher frequency of indeterminate vine growth habits and pigmented seed among photoperiod-sensitive accessions.

### Genome-wide association of descriptive traits

Due to the observed population structure and differences in LD between groups, and given the limited sample size available for Andean accessions, the focus of the genome-wide association studies conducted herein was on Mesoamerican accessions (*k1*, *k3*-*k5*, and *k8*). To detect descriptive trait-associated genetic variants in these Mesoamerican accessions (*n*=619), a set of 19,398 markers was utilized. Q-Q plots presented in Figure S6 indicated adequate model fit for the data. A total of 169 significant marker-trait associations were identified across the descriptive traits (Fig. 2, Table S6). Forty SNPs were detected at least twice by the four models (rTASSEL MLM, rMVP MLM, FarmCPU, and BLINK), and 12 SNPs were detected at least three times. The SNPs detected for each trait with multiple models are presented in Table S7 and described in the text. Twenty-two significant SNPs were detected for flower color; all 11 hits on chromosome Pl06 were within each other’s LD-defined search space. Photoperiod sensitivity, seed coat color, and determinacy had six, four, and three significant SNPs, respectively.

**Fig. 2.**
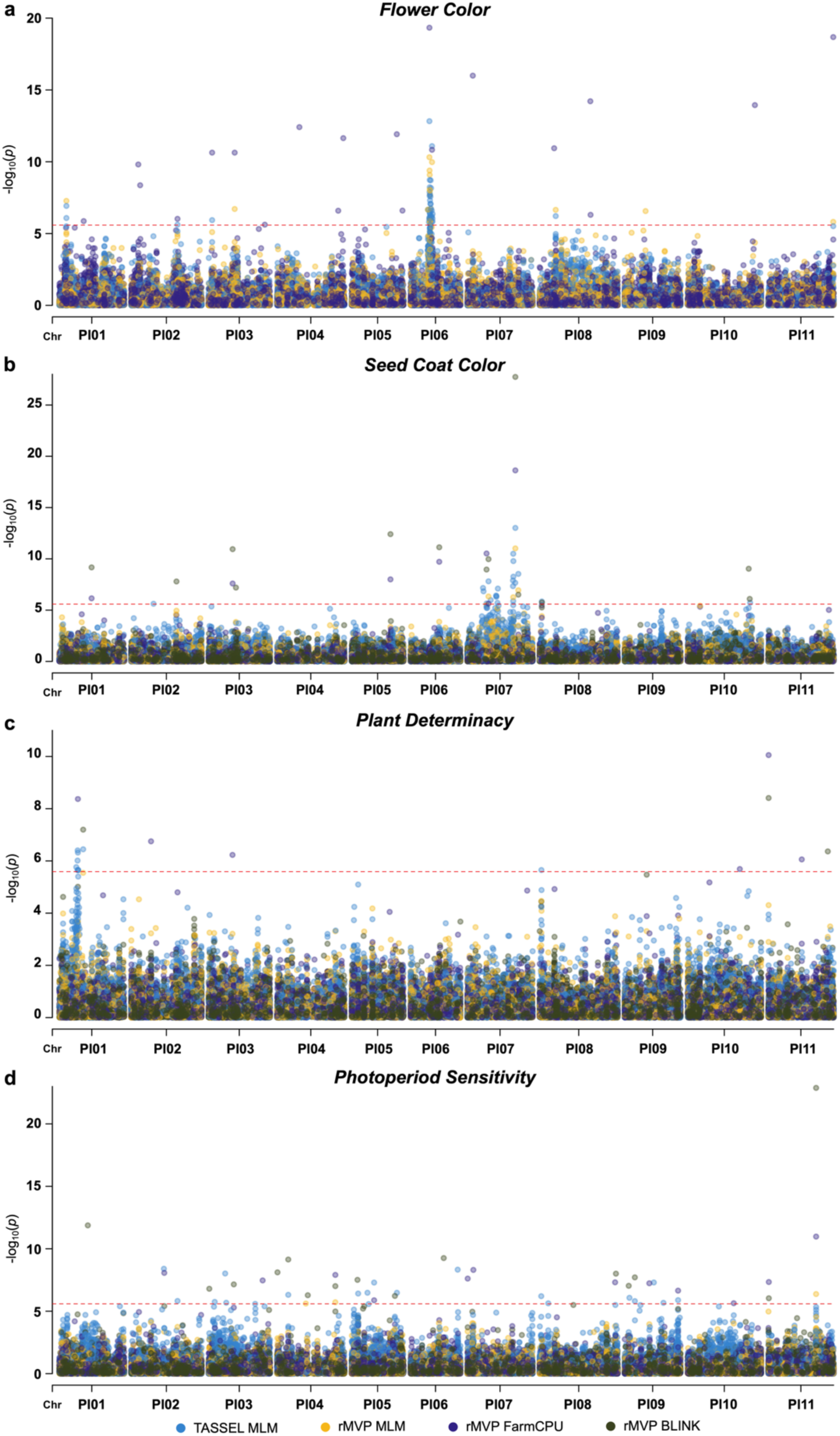
Genome-wide associations for descriptive traits. Models (TASSEL MLM, MVP MLM, MVP FarmCPU, and BLINK), indicated by color, were compared for **a)** flower color, **b)** seed coat color, **c)** determinacy, and **d)** photoperiod sensitivity among 619 Mesoamerican accessions. The vertical axis represents the significance of the association with the red dashed line designating the Bonferroni corrected significance threshold (⍺=0.05). The horizontal axis represents the chromosomal position of each single nucleotide polymorphism for the eleven lima bean chromosomes.

*Candidate genes for flower color, seed coat color, determinacy, and photoperiod sensitivity* Candidate genes were explored for the 40 SNPs detected by multiple models using a search space of 650 kb (i.e., ±325 kb). These genes were filtered based on homology with *P. vulgaris* and evidence of gene expression in relevant tissues in common bean (O’Rourke et al 2014). The most promising candidates can be found in Table S8. A total of 200 genes were filtered from a list of 500 to be potential candidates in the genomic regions significantly associated with flower color. Among these was the flavonoid 3′5′ hydroxylase (F3’5’H) gene (*Pl06G0000104100*), also known as the *V* gene in common bean, and dihydroflavonol-4-reductase (DFR; *Pl11G0000399000*), another key gene involved in flavonoid biosynthesis. Other potential candidate genes included transcription factors (TFs) involved in regulation of pigments, such as a WD repeat-containing (WDR) protein. A total of 142 genes were filtered from a list of 380 genes in genomic regions associated with seed coat color. Among these, a MYB TF and a glutathione S-transferase (GST) family protein were identified across multiple regions of Pl07.

A list of 232 genes were identified in genomic regions associated with determinacy, and 146 were considered for candidate genes. Among these, three candidates were identified for their roles in shoot apical meristem and gravitropism associated with plant architecture including a WUSCHEL-related gene (*Pl08G0000016500*), *shoot gravitropism 2* (*SGR2*; *Pl11G0000005800*), and a *MADS-box TF 6* (*Pl11G0000009100*).

Photoperiod sensitivity-associated genomic regions included a total of 558 genes. Among the filtered 315 genes in these regions, five relevant flowering-related candidate genes were identified including *Flowering Locus T* (*FT*) (*Pl04G0000305000*), *Histone lysine N-methyltransferase CONSTANS-like 2* (*CO*) (*Pl11G0000016000*), and *TEOSINTE BRANCHED 1/CYCLOIDEA/PCF 4-like* (*TCP4-like*) (*Pl11G0000293200*). The *MADS-box TF 6* (*Pl11G0000009100*) was in the search space for two significant SNPs (146 kb apart), one for each of determinacy and photoperiod sensitivity.

### Seed trait variation across environments

A partially overlapping subset of the panel (*n*=141–308, totaling 353 accessions) was field-evaluated for seed traits including macronutrients (protein, starch, fat, and ash) and HSW in six field environments (Central Ferry 2023 and 2024, CVARS 2023 and 2024, and Davis 2024 and 2025; Table S9). The number of accessions grown in common in each environment can be found in Figure S7. Macronutrients were predicted with high accuracy (*r*=0.73–0.99) via NIRS using a custom calibration that outperformed the pre-existing VPM calibration (Table S2). Across all samples, protein ranged from 13.1–34.0%, starch from 16.2–54.2%, fat from 0.4–1.8%, and ash from 4.2-6.0%.

A PCA of seed traits in the 353 field-evaluated accessions revealed differences among protein, starch, fat, and ash content across growing environments. PC1 explained 73.4% of the variation, with protein being significantly negatively correlated with starch both within and across environments (Fig. 3a, Figure S8). PC2 explained 17.3% of the variation and corresponded to differences between Mesoamerican (and admixed) vs. Andean accessions. Distributions of seed traits varied by subpopulation and environment (Fig. 3b). For example, mean protein content was higher in Davis and Central Ferry compared to CVARS across subpopulations, whereas mean fat content was lower in Davis and Central Ferry. The Andean accessions had higher HSW, protein, ash, and fat content and lower starch content in the Davis and Central Ferry environments compared to the Mesoamerican and admixed accessions.

**Fig. 3.**
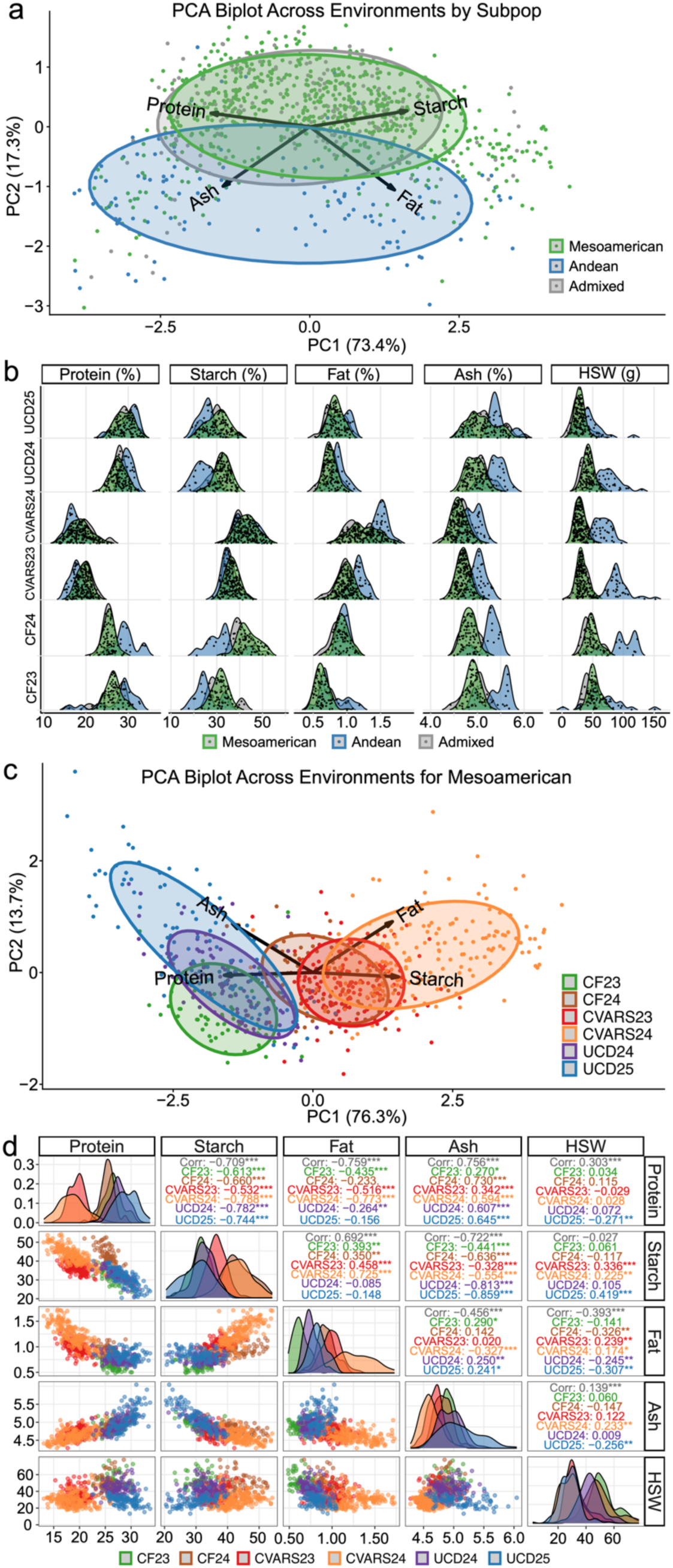
Seed trait analysis (macronutrient traits and hundred-seed weight (HSW)) for field-evaluated accessions. **a**) Principal component analysis (PCA) of macronutrient traits for the 353 accessions grown in any of the six environments (Central Ferry 2023 (CF23), Central Ferry 2024 (CF24), Coachella Valley 2023 (CVARS23), Coachella Valley 2024 (CVARS24), Davis 2024 (UCD24), Davis 2025 (UCD25)) grouped by subpopulations: Mesoamerican (*n*=248), Andean (*n*=47), and admixed (*n*=56). **b)** Density distributions of macronutrient traits (protein, starch, ash, fat) and HSW for the three subpopulations. **c)** PCA of macronutrient traits for 248 Mesoamerican accessions, colored by environment. **d)** Pearson correlations, histograms, and scatterplots of 248 Mesoamerican accessions across seed traits and environments.

**Fig. 4.**
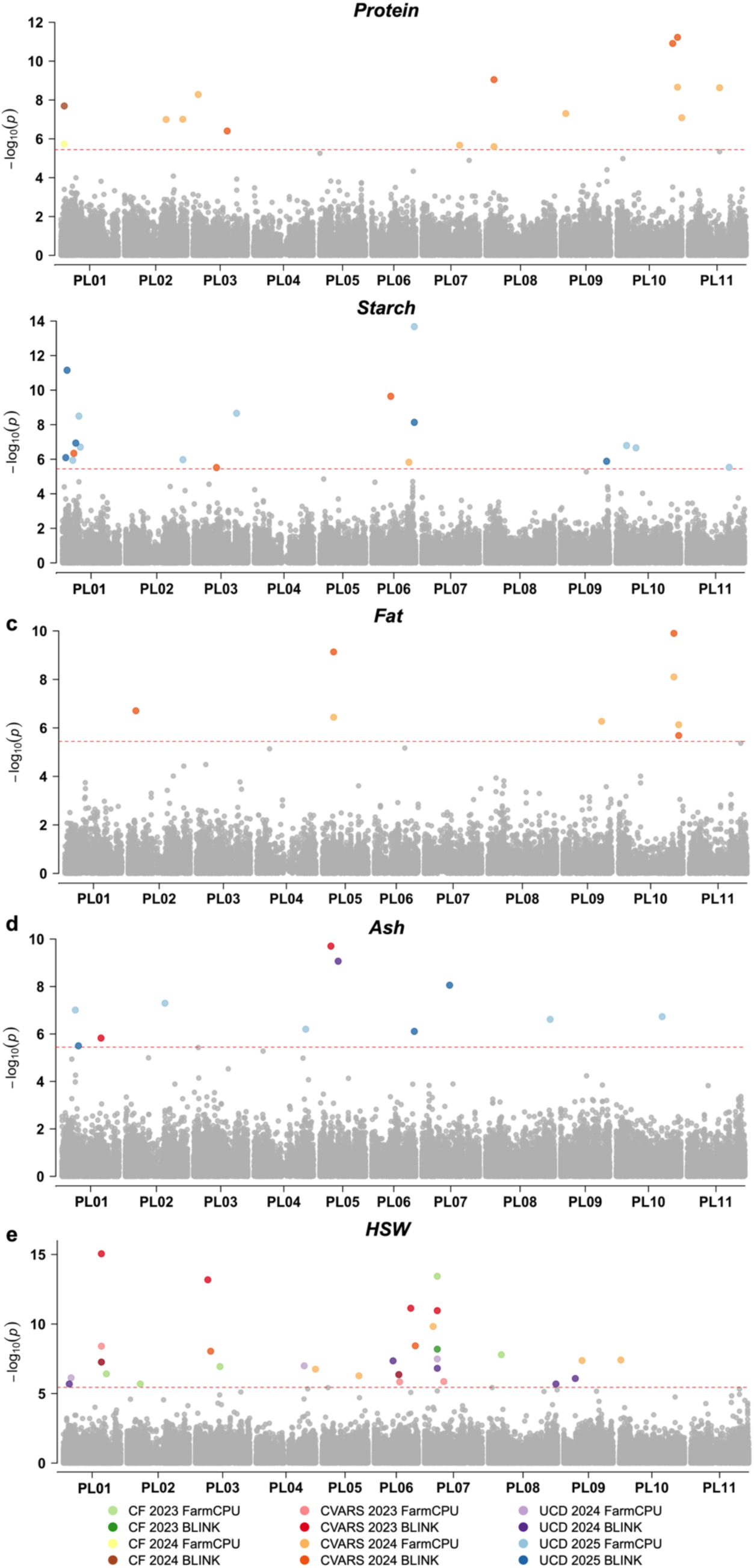
Genome-wide associations of seed traits in Mesoamerican accessions (*n*=248) for **a)** protein, **b)** starch, **c)** fat, **d)** ash, and **e)** hundred-seed weight (HSW) detected using multiple methods. Environmental abbreviations: Central Ferry 2023 (CF23); Central Ferry 2024 (CF24); Coachella Valley 2023 (CVARS23); Coachella Valley 2024 (CVARS24); Davis 2024 (UCD24); Davis 2025 (UCD25). The vertical axis represents the significance of the association with the red dashed line designating the Bonferroni corrected significance threshold (⍺=0.05). The horizontal axis represents the chromosomal position of each single nucleotide polymorphism for the eleven lima bean chromosomes. Colored points indicate significant associations for a given model and environment.

Due to the observed population structure, we focused our subsequent analyses on the 248 Mesoamerican accessions that were field-evaluated. A PCA of macronutrient traits within this subpopulation exhibited a similar pattern to the 353 accessions (Fig. 3c). PC1 explained 76.3% of the variance and corresponded to differences among growing environments. In particular, seed grown in both years at CVARS had significantly lower protein (*p*<0.001) and significantly higher (*p*<0.001) fat content than in any other environment. Composition of seed grown in Central Ferry showed variation between years, driven by starch and fat content, and was more similar in 2024 to seed produced in both years at CVARS.

Within environments in the Mesoamerican subpopulation (Fig. 3d), protein was significantly negatively correlated with starch (*r* = -0.709, *p*<0.05) and fat (*r* = -0.759, *p*<0.05) and positively correlated (*r* = 0.756, *p*<0.05) with ash. The magnitude of correlation for protein and fat was highest in CVARS 2024 (*r* = -0.796, *p*<0.05) and lowest in Davis 2025 (*r* = -0.181, *p*<0.05).

A mixed analysis of variance (ANOVA) revealed significant (*p*<0.01) main and interaction effects of genotype, location, and year on all traits for the total of 353 accessions evaluated and the 248 Mesoamerican accessions (Figure S9, Table S10). A mixed ANOVA was also performed for the two field trials (years) conducted within each location. The genotype by year interaction was not significant for HSW within any locations (Figure S9). Genotype explained a high proportion of variance for HSW in CVARS (72%) and Central Ferry (65%), but not Davis (19%), where 30% of the variance was explained by replicate within year. HSW showed high heritability within each location (*H*^2^=0.67 - 0.92, Table S10).

### Genome-wide association analyses for macronutrient and hundred-seed weight (HSW) traits

To detect seed trait-associated genetic variants in the Mesoamerican subpopulation (*n*=248), a set of 13,892 markers was utilized. Q-Q plots presented in Figure S10 showed that the models appeared to have a good fit for the data. A total of 94 unique significant marker-trait associations were identified across environments, methods, and traits (Table S11). Of these, 10 marker-trait associations were detected by two or more methods or across multiple environments (Table 3). Seven of these associations were identified in CVARS, while three and two were in Davis and Central Ferry, respectively. HSW was the only trait for which individual SNPs were detected in multiple environments, which was the case for each of Pl04:39931035 and Pl07:10553376. Two SNPs, Pl10:50347720 and Pl10:46341161, showed significant associations with both fat and protein in CVARS 2024 (Table 3 and Table S11). Two additional SNPs were detected across multiple traits by only one model: Pl01:11082760 for ash and protein in Davis 2025, and Pl01:11262701 for protein in Davis 2025 and starch in CVARS 2023.

**Table 3.**
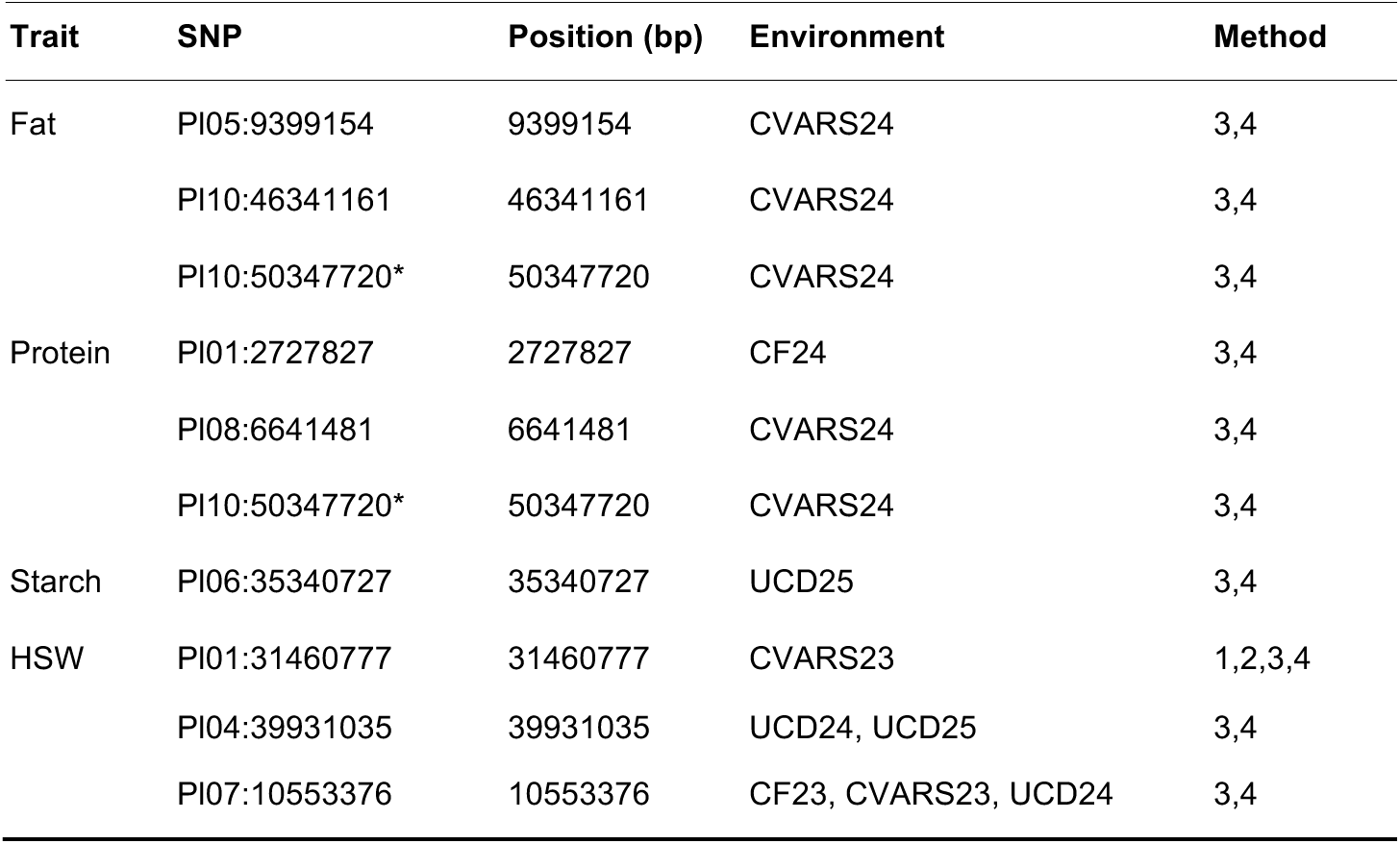
SNPs associated with seed traits in the field-evaluated Mesoamerican accessions, which were detected in at least two methods. GWAS methods denoted: 1, TASSEL MLM; 2, rMVP MLM; 3, rMVP FarmCPU; 4, BLINK. * SNP detected for multiple traits.

| Trait | SNP | Position (bp) | Environment | Method |
| --- | --- | --- | --- | --- |
| Fat | PI05:9399154 | 9399154 | CVARS24 | 3,4 |
|  | PI10:46341161 | 46341161 | CVARS24 | 3,4 |
|  | PI10:50347720* | 50347720 | CVARS24 | 3,4 |
| Protein | PI01:2727827 | 2727827 | CF24 | 3,4 |
|  | PI08:6641481 | 6641481 | CVARS24 | 3,4 |
|  | PI10:50347720* | 50347720 | CVARS24 | 3,4 |
| Starch | PI06:35340727 | 35340727 | UCD25 | 3,4 |
| HSW | PI01:31460777 | 31460777 | CVARS23 | 1,2,3,4 |
|  | PI04:39931035 | 39931035 | UCD24, UCD25 | 3,4 |
|  | PI07:10553376 | 10553376 | CF23, CVARS23, UCD24 | 3,4 |

### Candidate genes identified for protein, starch, and fat content and HSW

A space of 500 kb (i.e., ±250 kb) was searched for statistically significant SNPs that were detected in multiple models and/or environments. Table S12 contains a list of candidate genes. Genes within genomic regions associated with seed protein content were explored for functions such as storage proteins, nitrogen assimilation and remobilization, and regulatory genes controlling seed filling and seed storage. A total of 92 genes were located within the regions surrounding significant SNPs for protein content. No candidate genes were identified with known relevance to the trait within the regions examined. As such, the search region was expanded to ±500 kb, containing a total of 223 genes. Two genes annotated as subtilisin-like serine protease 2 (*Pl01G0000029500, Pl08G0000080800*) and a Bowman-Birk trypsin inhibitor (*Pl08G0000072900*) were identified.

Genes contributing to starch content were assessed based on relevance to functions associated with starch biosynthesis and degradation, sugar transport and partitioning, and carbon metabolism. A total of 78 genes were identified in the region on chromosome 6 identified by the GWAS. Because there were no known relevant genes in the search space, the region was extended as was done for protein. A SWEET sugar transporter was identified (*Pl06G0000332400*), along with three carbohydrate-active enzymes with GO terms associated with carbohydrate metabolic processes.

Genomic regions associated with seed fat content were explored for genes with functions relating to fatty acid synthesis, triacylglycerol (TAG) assembly, and regulatory TFs controlling seed oil. In total, 95 genes were identified in the regions surrounding significant associations with seed fat content. Among them, two primary candidate genes were identified: acyl-coenzyme A thioesterase-like protein (*Pl10G0000282500*; IPR006683), which catalyzes the hydrolysis of acyl-CoA esters into free fatty acids, and *acyl carrier protein 4* (*Pl10G0000318600*; IPR003231; GO:0006633 (fatty acid biosynthetic process)). However, only *Pl10G0000282500* has evidence of gene expression in common bean seed.

Genes related to hundred-seed weight were explored across a total of 80 genes located in three regions across chromosomes 1, 4, and 7. Genes were evaluated based on descriptions and functional annotations related to cell division, cell expansion, carbon accumulation, and hormones involved in growth such as auxin and cytokinin. We identified multiple candidate genes including two peptide transporters *NRT2/PTR Family 3.1-like* (*Pl01G0000252800*, *Pl01G0000252900*), two cyclin family proteins involved in cell division (*Pl01G0000253600*, *Pl01G0000253700*; IPR014400), *4-alpha-glucanotransferase DPE2-like* involved in starch metabolism (*Pl01G0000251000*), and a papain family cysteine protease (*Pl07G0000110800*) and serine carboxypeptidase (*Pl07G0000110600*) involved in seed storage proteolytic processing.

### Flowering time for day-neutral accessions grown in two environments

Flowering time was evaluated across the union of 353 accessions grown in Davis in 2024 and 2025 (Table S13). The distribution of flowering time for each subpopulation varied across years, with a wider distribution in 2024 compared to 2025 (Fig 5a). Bush types on average flowered earlier than vine types in both years (Fig 5b). Bush types flowered in similar timing across both 2024 and 2025, while vine types had later flowering in 2024. In a mixed ANOVA for flowering time, the effects of genotype, environment, and their interaction were all significant (*p*<0.001; Figure S11, Table S10).

**Fig. 5.**
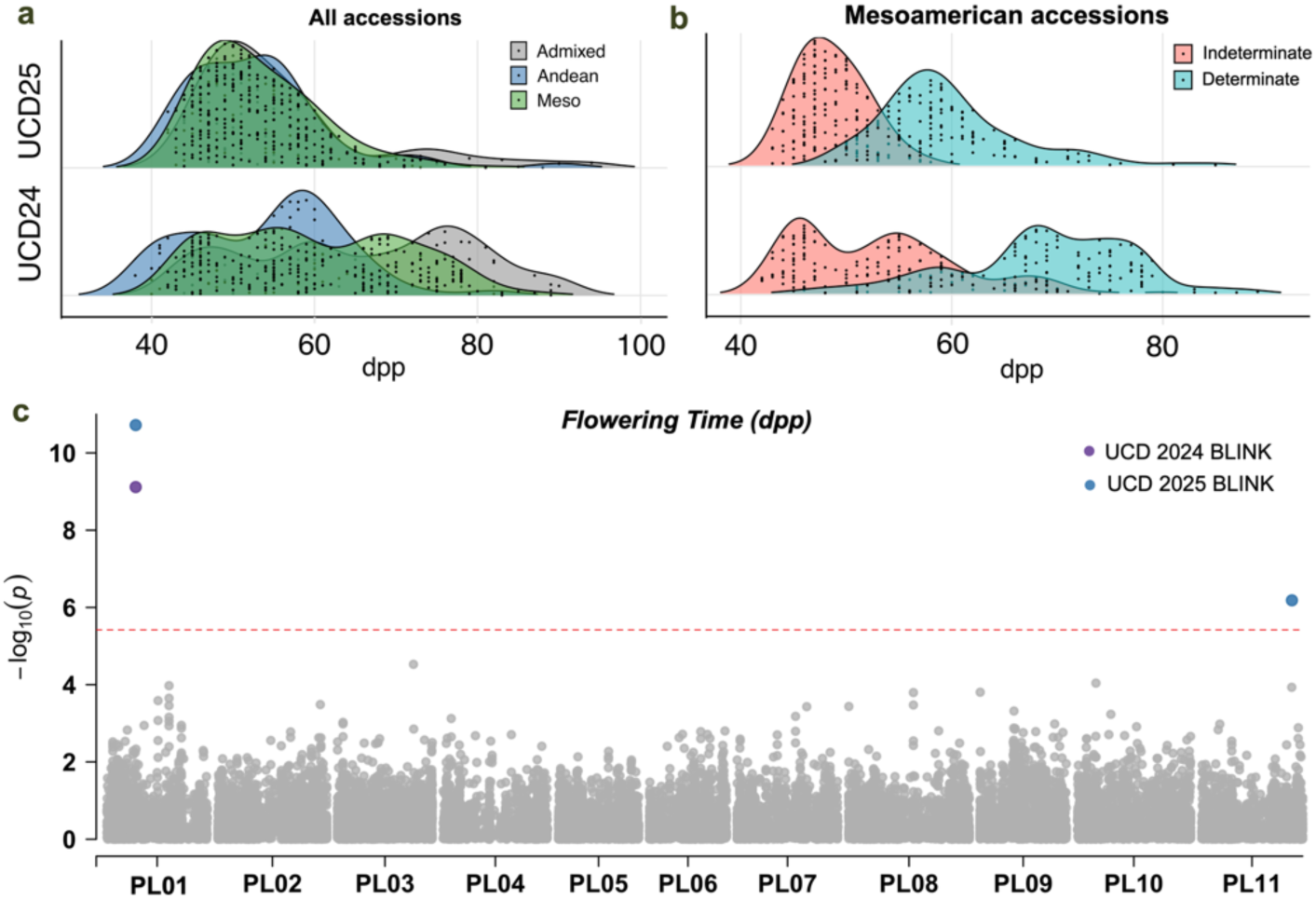
Flowering time analysis for accessions field-evaluated in Davis 2024 (UCD24) and 2025 (UCD25). Density distributions for a) all 353 accessions and b) 127 Mesoamerican accessions. c) Genome-wide associations in Mesoamerican accessions; colored points indicate significant associations detected with BLINK in a given environment.

A GWAS was performed for the Mesoamerican accessions grown in 2024 and 2025 in Davis (*n*=127). All four models were evaluated, and two significantly associated markers (Pl01:13602593 and Pl11:43014163) were identified with BLINK. The Q-Q plots (Figure S11) demonstrated a good fit for the data. Pl01:13602593 was detected in both years. Candidate genes were searched with use of gene annotations to identify genes involved in flowering. Two candidates were identified within a LD-defined search space of 700 kb (± 350 kb, Figure S2): a serine/threonine-protein kinase PBS1-like (*Pl01G0000122500*) involved in shoot apical meristem maintenance, and a trehalose-6-phosphate synthase (*Pl11G0000350000*) involved in signaling flowering time.

### Genomic prediction for descriptive, agronomic, and seed traits

Genomic prediction was conducted for descriptive traits for the 619 Mesoamerican accessions using the same datasets as in GWAS (Fig. 5a). Median predictive abilities ranged from 0.59– 0.78 across traits. Flower color (0.65–0.81), seed coat color (0.70–0.79), and determinacy (0.73–0.82) had among the highest predictive abilities. Photoperiod sensitivity displayed lower predictive ability ranging from 0.53 to 0.64.

Predictive abilities were also analyzed for seed traits for the 248 Mesoamerican subpopulation accessions using the same datasets as in GWAS. Median predictive abilities ranged across environments from 0.37 to 0.72 for ash, 0.15 to 0.70 for fat, 0.41 to 0.62 for protein, 0.25 to 0.66 for starch, and 0.39 to 0.80 for HSW depending on the environment (Fig. 6b). HSW had the highest predictive abilities across traits specifically in the Central Ferry 2023 and CVARS environments. Fat had lower predictive abilities compared to other traits, except in CVARS 2024 where median accuracy was 0.705. Among the six environments, ash had the highest predictive abilities in Davis while starch had the highest predictive abilities in CVARS and Davis. Flowering time among day-neutral accessions (*n*=127) grown in Davis demonstrated low predictive abilities in both years.

**Fig. 6.**
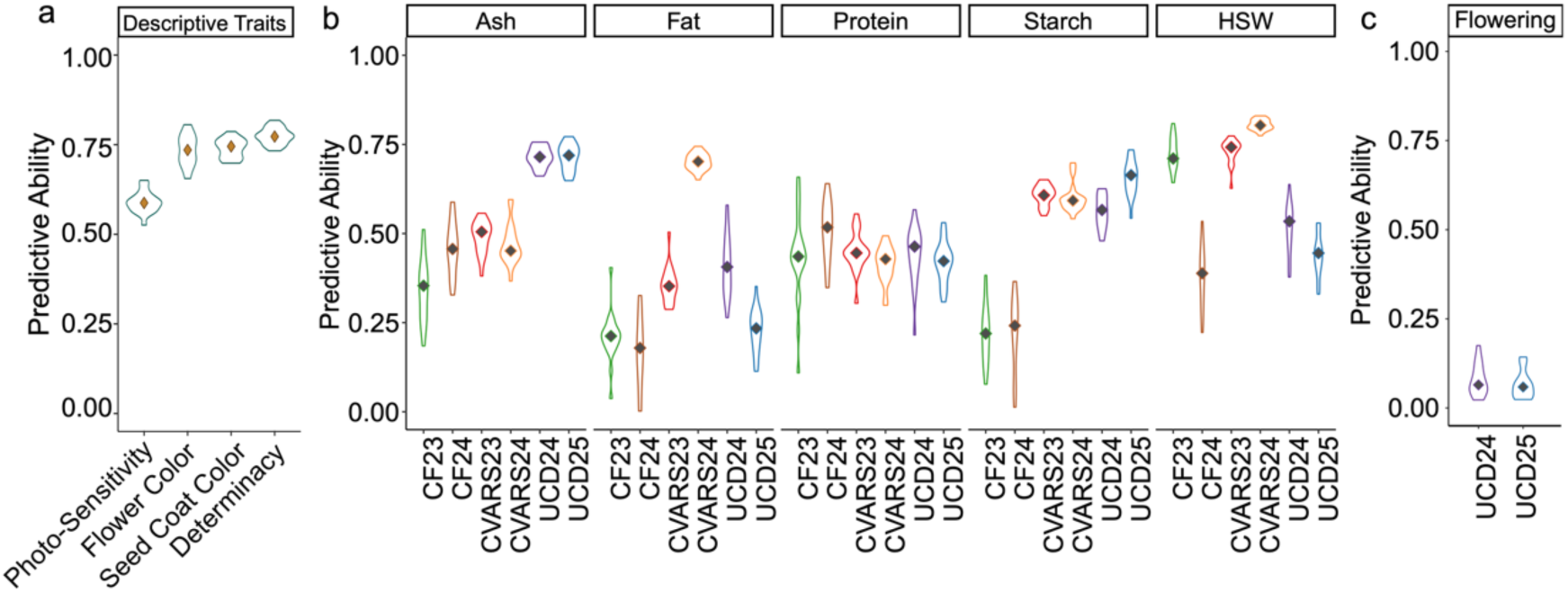
Genomic predictive abilities for **a)** descriptive traits (photoperiod sensitivity, flower color, seed coat color, and plant determinacy) with 619 accessions, **b)** seed traits (protein, fat, starch, ash, and hundred-seed weight (HSW)) with 248 accessions, and **c)** flowering time (in days post-planting) for 127 accessions. Seed traits and flowering time were predicted in corresponding field trials: Central Ferry 2023 (CF23), Central Ferry 2024 (CF24); Coachella Valley 2023 (CVARS23); Coachella Valley 2024 (CVARS24); Davis 2024 (UCD24); Davis 2025 (UCD25).

**Fig. 7.**
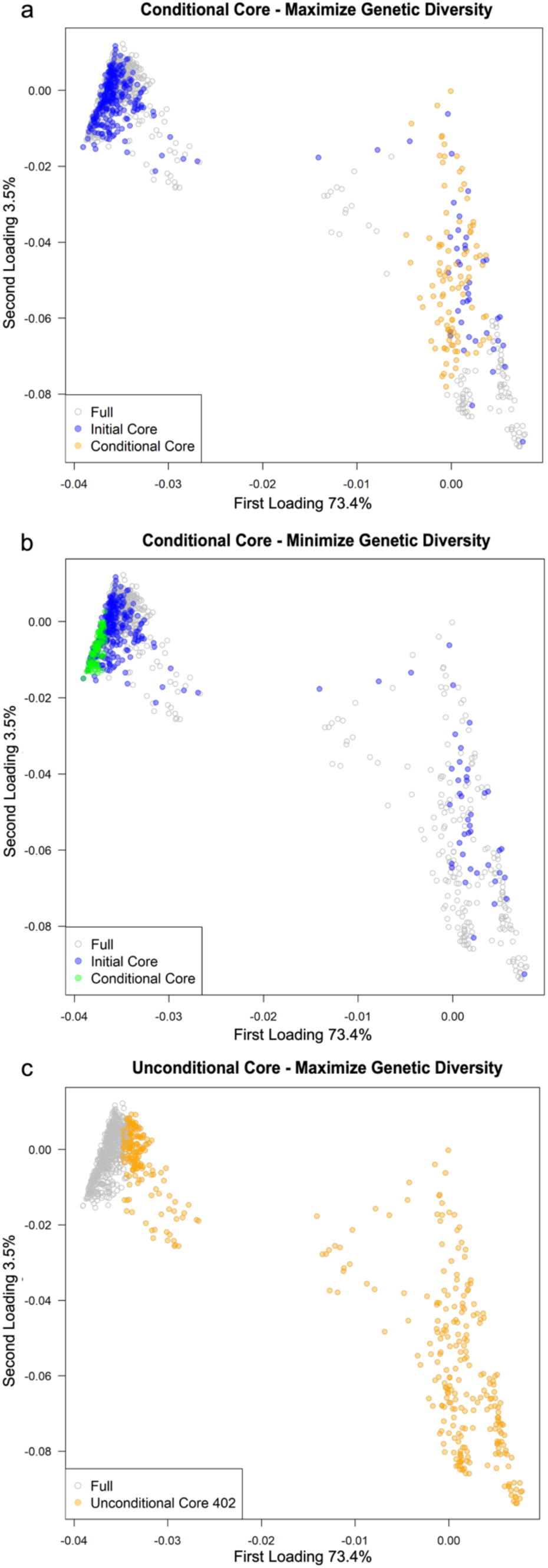
Results from use of a super-saturated method to select, conditional upon the inclusion of 305 samples (labeled ‘initial core’), a set of 100 supplemental samples (labeled as ‘conditional core’) that **a)** maximize and **b)** minimize genetic diversity; and to select **c)** an unconstrained core collection (*n* = 402). Within each panel, unselected samples are indicated via open grey circles.

### Conditional and Unconstrained Core Collections

Of the samples constrained to be in the conditional core, all 305 were domesticated, and 207 (including 150 from the US) were evaluated in at least one of the six field environments analyzed above. Of the 305 samples, 231 (including 122 in *k5*, 34 in *k2*, 21 in *k8*, and 41 admixed) were from the US, 20 from Brazil (of which 18 were in *k4*), 10 from Colombia (including 6 in *k4*), and 11 from Mexico (including 6 in *k3*). Counts ≤5 are not described in this section and can be found in Table S1. We selected a set of 100 supplemental samples that maximized genetic diversity (conditional upon the 305 samples being included) to form a ‘conditional core’ collection. The E(S^2^) value reached was 93,273.67, in 87 iterations. Of these 100 samples, 12 in *k2* were from the US, 11 in *k2* from Peru, 7 in *k2* from Ecuador, 11 (wild and in *k6*) from Costa Rica, 16 (wild and in *k6*) from Guatemala, and 15 in *k7* from Brazil. In total, 52 of these 100 samples were domesticated, 46 wild, and 2 unknown; and 64 were designated as photoperiod-sensitive, 33 day-neutral, and 3 unknown. Thirteen of the selected samples were evaluated in at least one of the six field environments. Finally, 18 pairs of selected samples were duplicates, meaning that those accessions were taken independently through single-seed descent and genotyped both at UC Davis and Clemson.

We also selected a set of 100 supplemental samples, alongside the 305 samples, that would minimize (rather than maximize) genetic diversity for purposes of a training set for use in genomic prediction/selection. The E(S^2^) value reached was 185,968.57, in 91 iterations. Of these 100 samples, 37 were from the U.S. (of which 28 were in *k5*) and 51 from Brazil (of which 45 were in *k4*), and the average HSW was 35.0 and 36.0 g, respectively. All 100 samples were domesticated, and 63 were designated as photoperiod-sensitive and 37 as day-neutral. None of the samples in this set were duplicates, and none were in the set selected for maximizing genetic diversity.

Finally, we selected an unconstrained core of 402 samples for comparison with the genetic diversity captured in the conditional core. The E(S^2^) value reached was 28,353.66, in 257 iterations. All 100 samples that were selected above to maximize genetic diversity were also selected for the unconstrained core. Of the 305 samples, 82 were selected for the unconstrained core, of which 64 were from the U.S. Within the 402 samples, 101 were from the US, 64 from Brazil, 51 from Peru, 36 from Mexico, 33 from Guatemala, 28 from Costa Rica, 26 from Colombia, 14 from El Salvador, 13 from Ecuador, and 8 from Nigeria. Of these, 326 were domesticated and 72 wild; and 268 were designated as photoperiod-sensitive, 129 day-neutral, and 5 unknown. A total of 122 pairs of selected samples were duplicates.

## Discussion

This study generated extensive phenotypic and genotypic data for a collection of diverse lima bean accessions primarily sourced from the USDA NPGS collection to assess population structure and identify the genetic basis of descriptive, agronomic, and seed traits. We identified common and unique associations across multiple traits and environments. Our results highlight new loci linked to genes that could influence these traits while also providing insight into relevant homologs previously identified in common bean, providing information for use in marker development. Further, we demonstrated that genomic prediction could be a useful tool in lima breeding for both descriptive and seed traits.

We identified accessions from Mesoamerican and Andean domestications in the collection through combined approaches of fastSTRUCTURE, PCA, and neighbor-joining tree analysis. The group *k2* defined from fastSTRUCTURE included five and three wild accessions from Ecuador and Peru, respectively, as well as domesticated accessions from Peru, Ecuador and the US, supporting the designation of this group as belonging to the Andean gene pool. *k1*, *k3*, *k4*, *k5*, and *k8* were closely related and contained domesticated accessions from Mexico and Central America, corresponding to a Mesoamerican gene pool. This could represent the MI gene pool originating from central-western Mexico, but notably more than half of the wild accessions from Mexico were designated as admixed. The Andean accessions had higher HSW than the Mesoamerican and admixed groups, as observed previously (Gutiérrez Salgado et al. 1995). In addition to these two subpopulations, a third subpopulation (*k6* and *k7*) emerged as a potential independent Mesoamerican gene pool. This group could correspond to MII due to the wild seed originating from Guatemala and Costa Rica characterizing *k6*. Interestingly, *k7* included a large proportion of accessions from Brazil, including several medium-seeded accessions; however, the NJ tree analysis showed that *k6* and *k7* were closely related to *k1* (Fig. S4), supporting a Mesoamerican origin. The relationship between *k6* and *k7* could indicate migration of seed from Central America to Brazil as observed in common bean (Gepts 1988; Kwak and Gepts 2009), while the inclusion of larger-seeded types in this group could indicate hybridization or an independent origin of larger seed size distinct from the Andean domestication.

LD was analyzed across the collection and within the three subpopulations. Garcia et al. (2021) demonstrated that in lima bean, LD decay is influenced by population structure and overall remains high (ranging from 0.3-0.4) over distances of 1 Mb, with higher LD in domesticated groups than wild. The LD decay of the total panel and subpopulations followed this pattern with the exception of the Andean subpopulation, which decayed to nearly 0 at 1 Mb. Interestingly, the group of *k6* and *k7* exhibited overall low *r*^2^, reaching 0.15 after approximately 50 kb and had a slight upward trajectory as physical distance increased. This pattern was influenced by *k6* (Figure S5), which comprised wild accessions and exhibited a very similar LD pattern to wild MI and MII accessions previously reported, suggesting high diversity.

The descriptive traits assessed here are among important domestication traits for lima bean. We identified a MYB2-like TF for seed coat color. In *Medicago* and *Arabidopsis*, MYB2-like (an R3-MYB) repressed proanthocyanidin biosynthesis by disrupting R2R3-MYB‒bHLH‒WDR (MBW) complex formation or activity (Dubos et al. 2008; Thévenin et al. 2012; Jun et al. 2015; Jiang et al. 2024). R2R3-MYB transcription factors are important regulators of seed coat color and patterning in *P. vulgaris* (Parker et al. 2024, McClean et al. 2024). Roy et al. (2025) found the *P. vulgaris MYB2-like* to be expressed in developing seed coats. The major-effect allele for white seeds (and flowers) in *P. vulgaris* is the *P* locus on Pv07, which encodes the MBW bHLH (McClean et al. 2018). While the candidates we identified were on Pl07, our significant markers were not proximal to the *P* homolog. For flower color we identified the *V* homolog (F3’5’H; McClean et al. 2022) and *DFR*, which acts one step downstream of *V* (Roy et al. 2025; Duwadi et al. 2018; Tapia et al. 2024) and was identified as a candidate for pod phenolics and blue/yellow pod color in snap bean (Myers et al. 2019; Celebioglu et al. 2023).

We identified candidates for photoperiod sensitivity including *FT*, a key gene in flowering initiation, and multiple co-regulators including TFs *CO-like* and *TCP4-like* characterized in *Arabidopsis* and other species (Kubota et al. 2017; Liu et al. 2021). FT is a signaling molecule that promotes the transition from vegetative to reproductive growth and whose homologs have been validated across plant genera including legumes *Medicago* and *P. vulgaris* (Kwak and Gepts 2009; Laurie et al. 2011; Wickland and Hanzawa 2015). For photoperiod sensitivity and determinacy, we identified a MADS-box 6 with homology to *AGAMOUS*-like MADS-box protein AGL12 in *P. vulgaris.* Tapia-López et al. (2008) found *Arabidopsis xal1* (renamed from *AGL12*) mutants flowered late under long days, with *FT* downregulated and pleiotropy for root traits. Ectopic expression of grape *AGL12* in *Arabidopsis* promoted flowering among other traits (Mao et al. 2023). The homolog of *PHYA* on Pv01, identified for photoperiod sensitivity in *P. vulgaris* (Weller et al. 2019), was absent in our results. We identified a significant association situated 26 Mb from this homolog, in a similar location to a previously identified lima QTL for flowering time (Garcia et al. 2021). For flowering time in day-neutral accessions, a significant association was identified in both field evaluations on Pl01, which was 3 Mb from our hit for photoperiod sensitivity. The lack of significantly associated markers proximal to previously characterized *P. vulgaris* homologs such as *P* for seed coat color, *PhyA* for photoperiod sensitivity, and *fin*-*TFL1y* for determinacy (Kwak et al. 2008; Repinski et al. 2012; Kwak et al. 2012; Denning-James et al. 2025) could be due to insufficient marker density or low minor allele frequency (e.g., in instances of lineage-specific allelic variation, which has been observed for both *P* and *TFL1y* in common bean; McClean et al. 2018; Kwak et al. 2012). Studies with whole-genome sequencing data could further investigate those genes and the genomic loci identified herein in lima. Fine mapping through recent recombination and/or RNA-seq would further inform candidate gene identification and regulatory networks for descriptive, agronomic, and seed traits.

We identified multiple significant associations with seed protein, starch, fat, and ash content. Three SNPs had significant associations for multiple traits (protein with each of starch, fat, and ash). The role of pleiotropy versus linkage in these relationships could be examined further as demonstrated in cassava (Villwock et al. 2025). We identified two candidate genes for HSW, both nitrate/peptide transporters encoding NRT1/PTR FAMILY 3.1-like proteins. These proteins have been demonstrated to affect thousand-grain weight and increase nitrogen use efficiency in rice (*Oryza sativa*; Yang et al. 2023). We also identified a papain family cysteine protease for HSW. Papain family cysteine proteases are major endopeptidases in cotyledons and have been examined in *P. vulgaris*, *Glycine max*, and *Vicia sativa* and *V. narbonensis*, among other species (Zakharov et al. 2004; Esteban-García et al. 2010; Matsuoka & Maruyama 2026; Fischer et al. 2000; Müntz 1996). Those studies indicate roles in partial hydrolysis (and thus remobilization) of storage proteins during seed germination, filling, and/or maturation. Exopeptidases such as the serine carboxypeptidase identified for HSW (in the same search space as the papain family cysteine protease) have also been implicated in seed germination and filling, and in cereal seed storage protein hydrolysis (Tan-Wilson and Wilson 2011; Vorster et al. 2023). Transcriptomic, proteomic, and peptidomic profiling spanning grain development and subsequent germination would be informative in clarifying the roles of nitrogen-related candidate genes identified herein for HSW (nitrate/peptide transporters, a papain family cysteine protease, and a serine carboxypeptidase) and for protein (subtilisin-like serine proteases and a Bowman-Birk serine protease inhibitor; BBI) (Bera et al. 2023; Vorster et al. 2023). Serine proteases (classified as subtilisin-like or trypsin/chymotrypsin-like) are also major endopeptidases in developing seed. However, D’Erfurth et al. (2012) studied an *M. truncatula* mutant for an endosperm-localized subtilisin-like serine protease found to control seed size and found no effect on protein content. Finally, BBIs are one of two main classes of protease inhibitors in legume seeds and can inhibit both trypsin and chymotrypsin (Bera et al. 2023; Vorster et al. 2023). The crystal structure of a lima BBI has been well-characterized (Stevens et al. 1974; Debreczeni et al. 2003; Ahmad et al. 2023) with homology to the candidate identified herein. In *G. max*, Wang et al. (2025) conducted CRISPR/Cas9-mediated knockouts and Kim et al. (2025) conducted RNAi-mediated knockdowns of seed-specific BBIs, and both found seed protein content to be unaltered. We recommend continued high-throughput phenotyping and/or GP for HSW and seed protein content, with monitoring for co-variability with yield, while potential candidates undergo further examination.

HSW had the highest heritability in this study, consistent with high heritabilities observed for HSW in other *Phaseolus* spp. (Nienhuis and Singh 1988; White et al. 1994; Okii et al. 2018). While the genotype-by-environment interaction was significant among the accessions grown in multiple environments for HSW, substantial variance was explained by genotype within the CVARS and Central Ferry locations. Variation between replicates in Davis trials explained a large proportion of variance, likely due to pest pressure in 2025 that negatively impacted seed quality. Davis experienced two heat waves during flowering in 2024, which could underlie differences in seed traits and/or flowering time between years. The nutrient-poor sandy soils at CVARS may underlie the lower seed protein content observed there. Andean accessions across Davis and Central Ferry environments were notably higher in protein and lower in starch than Mesoamerican accessions, suggesting their larger seed size was not simply due to an increased proportion of starch. Islam et al. (2002) found Andean beans had higher phaseolin and lower lectin concentration than Mesoamerican beans in the CIAT common bean core collection. More detailed characterization of protein and carbohydrate composition in limas would be informative; the latter (e.g., via Couture et al. 2024) would also provide insight into dietary fiber. More spatial and temporal resolution on the accumulation of macro- and micronutrients in seed could elucidate relationships between seed physical and compositional traits. Anti-nutritional factors, such as cyanogenic glucosides (Montero-Rojas et al. 2013) and phytate (Asif et al. 2022), also need to be considered in breeding of lima beans.

Genomic predictive (GP) abilities were fairly high for most of the descriptive, agronomic, and seed traits studied herein. This finding suggests GP could be useful if already assaying limas on a low-to mid-density marker platform that is sufficient for capturing relatedness, as found to be the case for 500 markers in the southern U.S. rice gene pool (Cerioli et al. 2022). However, we would recommend keeping sufficient density to conduct genetic mapping as there are still priority traits and populations to be mapped. We would also recommend pooled genotyping (across multiple seeds of an accession) to capture the heterogeneity that can be characteristic of accessions stored in gene banks, as evidenced in the duplicates assayed herein and the number of duplicates from which both samples were selected for the unconstrained core.

A further step would be to examine the relatedness and complementarity of multiple lima germplasm collections through collaborations that pair genotypic (ideally on the same platform) and on-site phenotypic characterizations. Lima germplasm collections are maintained at CIAT, Cali, Colombia (Debouck et al. 2021); the International Institute of Tropical Agriculture, Ibadan, Nigeria (Temegne et al. 2024); the University of Puerto Rico, Mayagüez (Montero-Rojas et al. 2013); and other universities and national programs, e.g., in Brazil (Oliveira-Silva et al. 2017; Oliveira-Silva et al. 2019; da Costa Almeida et al. 2025). Several of these collections, alongside the USDA NPGS lima collection, are cataloged in Genesys (https://www.genesys-pgr.org/). On-site characterizations would mitigate the challenge of evaluating accessions with diverse geographic origins and, somewhat accordingly, distinct photoperiod requirements. Incorporating more wild accessions in characterizations, whether from other germplasm collections and/or from the unavailable inventory at USDA NPGS, would provide more resolution regarding domestication histories. Seed multiplication was challenging for the panel characterized herein; 810 accessions were genotyped, whereas only 353 had sufficient seed for field evaluation. Regenerating, maintaining, and distributing stocks remains a critical but challenging role of the USDA NPGS (Byrne et al. 2018). Converting more of the genotyped accessions to photoperiod insensitivity (as has been done in sorghum; Stephens et al. 1967; Rosenow et al. 2003; Winans et al. 2023; Patil et al. 2023; Crozier et al. 2024) would facilitate field evaluation and seed multiplication in the US. The 118 photoperiod-sensitive, domesticated accessions in the conditional core could be prioritized for conversion to form an optimized panel for mapping of useful traits and alleles. Further training set optimization for genomic prediction/selection could also be fruitful, as 31 samples from the U.S. and 57 samples from Brazil were selected to minimize genetic diversity alongside the samples constrained to be in the core from those countries, suggesting relatedness (at least within each of *k5* and *k4*, respectively) that could be useful in prediction. Finally, addition of collected genotypic and phenotypic data, using standard formats and definitions, into LIS, USDA-ARS GRIN, and other databases when pertinent, is important for continued characterization and utilization (Shrestha et al. 2012; Byrne et al. 2018; Arnaud et al. 2020; Volk et al. 2021).

In conclusion, this study provides a comprehensive genotypic and phenotypic evaluation of key descriptive, agronomic, and seed traits for a diverse panel of lima bean. Our results identify multiple loci and candidate genes underlying these traits and provide a foundation for applying genomic tools for holistic improvement of this crop.

## Acknowledgments

We gratefully acknowledge the undergraduate students who contributed to laboratory-based phenotyping including Maja Watte, Shunmitha Babu, and Rajdeep Bains. We gratefully acknowledge Martha Delgado for her assistance in seed processing and phenotyping. We gratefully acknowledge the field teams at the UC Riverside Coachella Valley Agricultural Research Station, UC Davis field station, and USDA Central Ferry Farm for their assistance in the field trials. We gratefully acknowledge Tyler Williams for greenhouse plant management and tissue sampling at Clemson. We also gratefully acknowledge Jonathan Berlingeri, Mary-Francis LaPorte, Sassoum Lo, and Tayah Bolt for their contributions to the development of the NIRS custom calibration script.

## Author Contributions

**Jaclyn A. Adaskaveg** coordinated field trials including seed stocks and experimental planning and logistics; curated accession metadata; was involved in quality control of genotypic data; coordinated collection of field- and laboratory-based phenotypes; developed and deployed the custom NIRS calibration; coordinated the conditional and unconstrained core analyses; conducted analyses of genotypic data (including population structure and LD), phenotypic data, and their integration, including genetic mapping and genomic prediction; prepared the figures and tables; and had a primary role in writing and revising the original draft.

**Jenna Hershberger** curated accession metadata, coordinated the genotyping effort at Clemson University, provided feedback on the conditional and unconstrained core experimental design, and was involved in funding acquisition.

**Andrew Farmer** coordinated the provision of genotypic data and uploading of that data to LIS, provided feedback in the quality control process for genotypic data, and was involved in funding acquisition.

**R. Varma Penmetsa** was involved in the genotyping effort at UC Davis and in the quality control process for genotypic data.

**Ivan Garcia-Lopez** contributed to field- and laboratory-based phenotyping.

**Julian Garcia-Abadillo** carried out the conditional and unconstrained core analysis.

**Xiangming Zhou** carried out the conditional and unconstrained core analysis.

**Bao-Lam Huynh** helped to facilitate field trials at CVARS.

**Philip Roberts** helped to facilitate field trials at CVARS and was involved in funding acquisition.

**Emmalea Ernest** provided feedback on the conditional and unconstrained core experimental design and was involved in funding acquisition.

**Marilyn Warburton** facilitated provision of USDA NPGS materials for use in this study and was involved in funding acquisition.

**Diego Jarquin** coordinated the conditional and unconstrained core analyses.

**Sarah Dohle** provided USDA NPGS materials for use in this study, guided the collation and interpretation of accession metadata, coordinated field trials at the USDA Central Ferry Farm, and was involved in funding acquisition.

**Antonia Palkovic** coordinated field trials including seed stocks and experimental planning and logistics, coordinated greenhouse-based grow-outs for genotyping and seed multiplication, and was involved in the genotyping effort at UC Davis and in funding acquisition.

**Travis A. Parker** coordinated the genotyping based at UC Davis, which included the ordering (via USDA-ARS GRIN) and collation of seed stocks and coordination of the greenhouse-based growout including sampling for genotyping and observation of descriptive traits; coordinated the provision of genotypic data; was involved in the quality control process for genotypic data; conducted preliminary GWAS for descriptive traits; and was involved in funding acquisition.

**Paul Gepts** designed and directed the USDA-NIFA-SCRI-funded project, was involved in the quality control process for genotypic data, provided feedback on population structure results and interpretation, participated in the revision of the original draft, and led the funding acquisition.

**Christine H. Diepenbrock** coordinated field trials including experimental planning and logistics; was involved in the quality control process for genotypic data and custom NIRS calibration development; guided analysis of genotypic and field- and laboratory-based phenotypic data including LD analysis, GWAS, and GP, and conditional and unconstrained core analyses; had a role in writing and revision of the original draft; and was involved in funding acquisition.

**All authors** critically reviewed the manuscript and approved it for submission.

## Data Availability

The raw sequence and genotypic data generated in this study will be made available at NCBI Bioproject XXX at time of manuscript acceptance and is being integrated into LIS, respectively. The phenotypic data generated in this study can be found in the supplemental materials and is being integrated into the USDA-ARS GRIN database. Custom scripts used in the analyses conducted in this study are provided as supplemental materials for the manuscript peer review process and will be available through a public GitHub repository for which a digital object identifier will be generated via Zenodo at time of manuscript acceptance.

