## Supplemental Figures for "Genotypic and multi-environment phenotypic evaluation of the lima bean USDA National Plant Germplasm System collection"

### **Figure S1.**

Summary statistics on unfiltered SNP data (134,309 SNPs): **a)** SNP density across the 11 chromosomes of *P. lunatus* colored by density of SNPs. **b)** Minor allele frequency, **c)** missingness by site, and **d)** heterozygous proportion by site with cutoffs indicated with vertical dashed line.

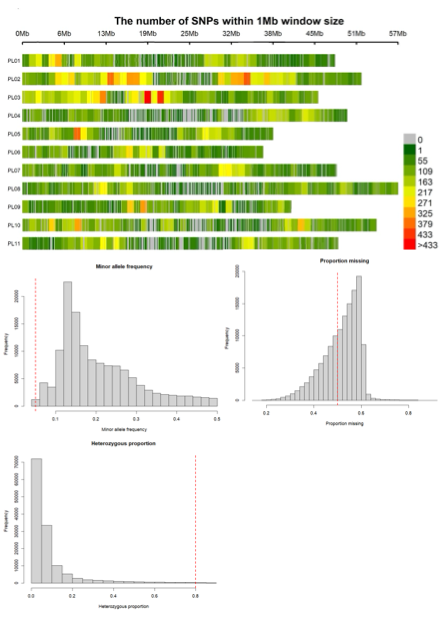

### Figure S2.

Linkage disequilibrium decay for GWAS datasets for the descriptive (a,b), multi-environment seed traits (c,d), and multi-environment flowering time (e,f) trait datasets. All accessions with phenotypic data (a, c, e) were analyzed as well as the Mesoamerican accessions in the dataset (b, d, f).

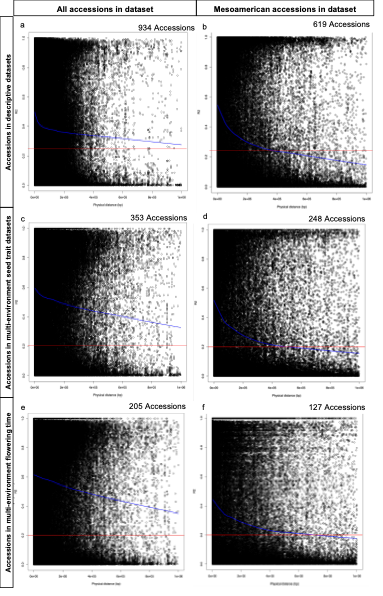

### **Figure S3.**

PCA of genotypic data with accession duplicates. Accession duplicates (taken independently through SSD and genotyped both at UC Davis and Clemson University) are indicated with a black line drawn between each duplicate.

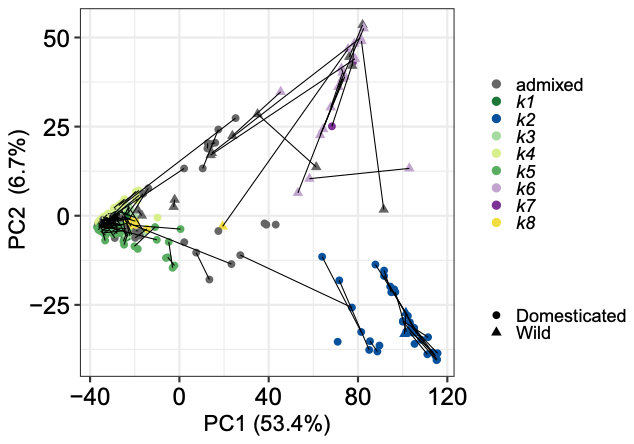

### **Figure S4.**

PCA of genotypic data colored by a) photoperiod sensitivity, b) HSW, and c) continent, d) PCA plotting PC1 and PC3 colored by fastSTRUCTURE *k* group at K=8, e) neighbor-joining tree colored by *k* group at K=8.

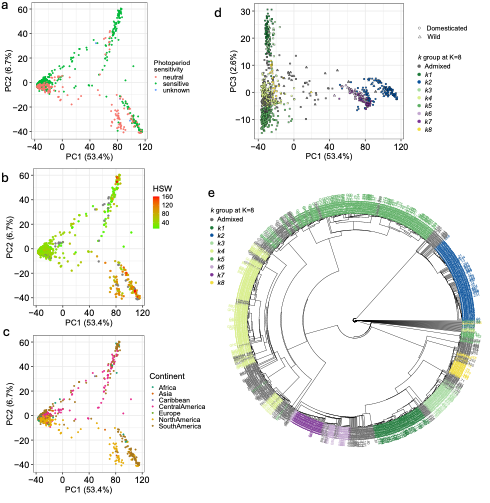

### **Figure S5.**

Linkage disequilibrium decay plots by individual fastSTRUCTURE *k* group at K=8. a) *k1* (n=50), b) *k2* (n=127), c) *k3* (n=50), d) *k4* (n=202), e) *k5* (n=264), f) *k6* (n=40), g) *k7* (n=46), h) *k8* (n=28). Red line indicates R^2^ of 0.2.

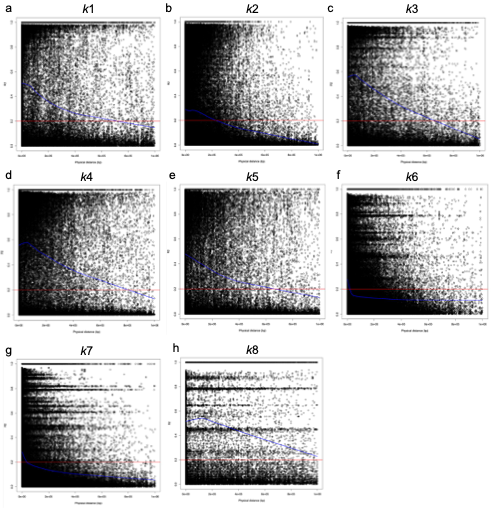

### **Figure S6**.

Q-Q plots for expected vs observed log_10_ of p-value for descriptive traits (photoperiod sensitivity, flower color, seed coat color, and determinacy) for GWAS with Mesoamerican accessions (n=619). Colored dots indicate methods used (TASSEL MLM, rMVP MLM, rMVP FarmCPU, and BLINK).

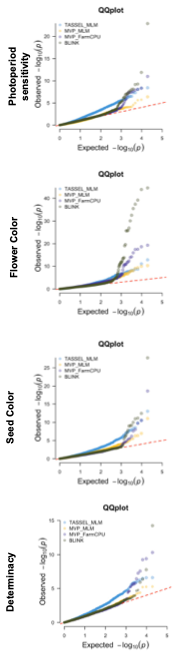

### **Figure S7**.

The number of accessions that were overlapping (i.e., evaluated in common) in each pairwise combination of environments. The number in parentheses under each environment name, within the axis labels, indicates the total number of accessions phenotyped in that environment. Darker yellow shading corresponds with a higher number, and lighter yellow with a lower number.

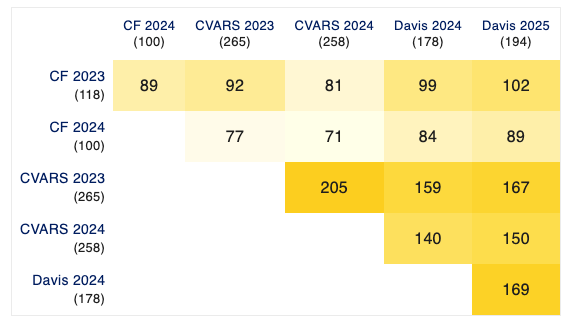

### **Figure S8**.

**a)** Principal component analysis (PCA) of macronutrient traits for the 353 accessions grown in any of the six environments (Central Ferry 2023 (CF23), Central Ferry 2024 (CF24), Coachella Valley 2023 (CVARS23), Coachella Valley 2024 (CVARS24), Davis 2024 (UCD24), Davis 2025 (UCD25)) grouped by field experiment. **b)** Pearson correlations, histograms, and scatterplots of 353 accessions grown across seed traits and environments. Significance is indicated by asterisks where *** <0.001, ** <0.01, * <0.05, and no asterisk indicates no significance.

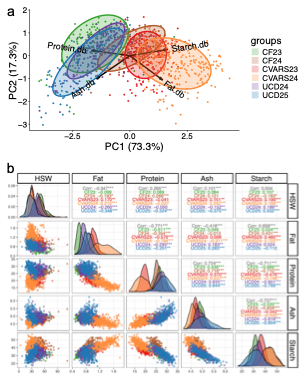

### **Figure S9.**

Variance components percentage from mixed model analysis across six environments for a) 353 field-evaluated, b) 248 Mesoamerican (Meso) accessions, c) 47 Andean accessions, d) 56 admixed accessions and two years within each location e) Davis, CA, f) Coachella Valley Agriculture Research Station (CVARS), CA g) Central Ferry, WA.

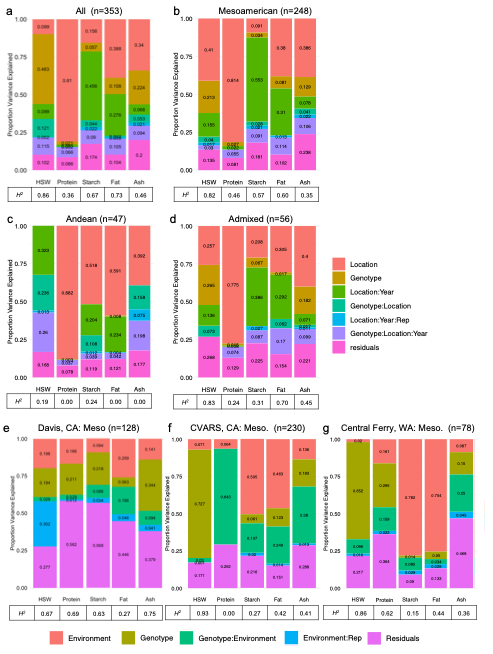

### **Figure S10.**

Q-Q plots for seed traits (HSW, protein, fat, ash, starch content) for GWAS analysis conducted in each environment (Central Ferry 2023 (CF23), Central Ferry 2024 (CF24), Coachella Valley 2023 (CVARS23), Coachella Valley 2024 (CVARS24), Davis 2024 (UCD24), Davis 2025 (UCD25)). Each plot is colored by model tested (TASSEL MLM, rMVP MLM, rMVP FarmCPU, and BLINK).

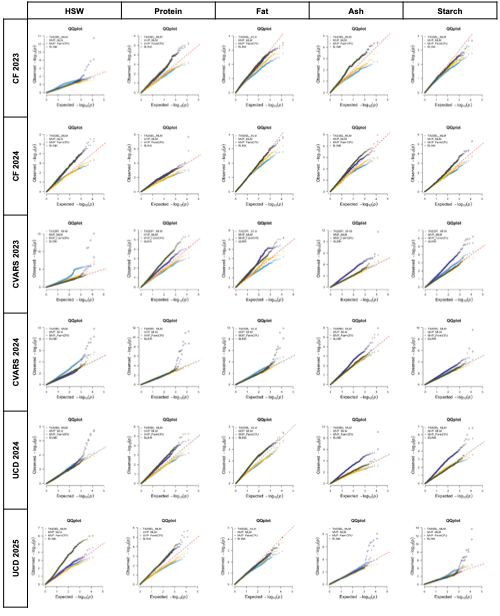

### **Figure S11.**

Variance components percentage from mixed model analysis for the multi-environment flowering time dataset in a) all evaluated accessions and b) Mesoamerican accessions. c) Q-Q plot for flowering time (days post planting, dpp) in Davis 2024 and 2025 where each plot is colored by model tested (TASSEL MLM, rMVP MLM, rMVP FarmCPU, and BLINK).

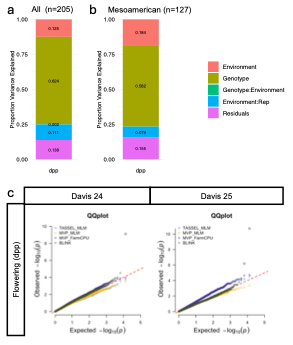
